# Machine learning general transcriptional predictors of plant disease

**DOI:** 10.1101/2023.08.30.555529

**Authors:** Jayson Sia, Wei Zhang, Mingxi Cheng, Paul Bogdan, David E. Cook

## Abstract

Plants utilize an innate immune system to defend against all classes of microbial invaders. While we understand specific genetic determinants of host-pathogen interactions, it remains less clear how generalized the immune response is to diverse pathogens. Using a data-driven approach, and utilizing feature selection based on network science and topology, we developed machine learning models that could predict host disease development across diverse pathosystems. These machine learning models identified early transcriptional responses predictive of later disease development, regardless of pathogen class, using a fraction of the host transcriptome. The identified gene sets were not enriched for canonical defense genes, but where statistically enriched for genes previously identified from independent data sets, including those described as representing a general plant stress response. These results highlight novel components of a general plant immune response, and demonstrate the application of machine learning to address biological hypotheses of a complex multigenic outcome.

**Teaser:** A machine learning approach can predict plant disease development caused by diverse microbial invaders, and newly identified genes may represent novel components of a general plant response to infection.

## Introduction

Plants rely on an innate immune system to counter infection. A hallmark of the plant innate immune system is the detection of invasion through diverse receptor catalogues that survey both the apoplastic and cytoplasmic compartments [1,2]. At the cell surface, membrane anchored pattern recognition receptors (PRRs), in the form of receptor-like kinases and receptor-like proteins, have been well described for their ability to detect a wide variety of ligands to initiate immune responses [3–5]. Perception of invasion at the cell-surface can trigger a number of cellular responses, including ion flux, reactive oxygen species production, post-translational modifications, and transcriptional reprograming [6,7]. These events control a myriad of physical and chemical defenses that collectively contribute to plant defense. Intracellular receptors, referred to as nucleotide-binding and leucine-rich repeat receptor (NLR) domain-containing proteins, detect non-self and modified-self ligands to initiate a plant defense response [8,9]. Structural data shows that different types of activated NLRs form multimer protein complexes, such as the pentamer ZAR1-RKS1 (HopZ-activated resistance 1, resistance-related kinase 1 [10], and tetramers ROQ1 (recognition of XopQ1) [11], and RPP1 (recognition of *Peronospora parasitica* 1) [12]. Collectively, the extracellular and intracellular immune receptors form a coordinated immune response to integrate detection, signal transduction, and response to diverse biotic incursions [13–16].

Plant immune receptors detect specific immunogenic ligands of diverse origins, but it is less clear how plants integrate diverse signals to achieve specific immune responses [17]. An open question concerns the extent to which immune responses are fine-tuned to specific ligands that enact a tailored defense response to the invader [13,18]. Analysis of plant transcriptional responses to diverse pathogens and ligands suggests there is both overlap and divergence to differing immunogenic signals. For instance, *Arabidopsis thaliana* seedlings responding to either a plant cell wall derived oligogalacturonides or bacterial derived flagellin peptide, elicit similar early transcriptional responses that diverged with time [19]. Also in *A. thaliana*, early transcriptional responses were largely overlapping in response to seven diverse immunogenic ligands, but also, the FLS2-flg22 interaction had a substantial number of unique transcriptional responses [18]. Early transcriptional response of *A. thaliana* to genetically diverse *Botrytis cinerea* isolates showed that wild-type Col-0 ecotype displayed significant transcriptional variation for a number of transcripts involved in hormone related signaling and PRR responses [20]. Despite the diverse transcriptional responses to the genetically diverse pathogen isolates, final disease development outcomes are rather limited [20], which can be conceptualized as many roads leading to the same place. Such a buffered immune response to diverse inputs may be mediated by interconnected immune subnetworks, providing robustness to the diversity of microbial interactors [13,21,22]. It is clear that activation of different PRRs do not provide identical transcriptional outputs [23], but immune signaling can channel responses from diverse inputs into largely overlapping outputs [13,20,18]. The emerging systems view of plant immunity suggests a highly interconnected network involving receptor detection, intracellular signaling, hormone cross-talk and cellular outputs to provide immunity, recently reviewed [17,24–27].

Machine learning (ML) represents a broad class of algorithms designed to optimize a function that relates input to output data. Highly predictive ML algorithms, such as random forests [28] and support vector machines [29] are well developed, and require less data abstraction through hidden states, making their output and learning human discernable. This contrasts deep learning techniques of the last decade that rely on highly-abstracted feature selection techniques, able to predict the most complex interactions, but obfuscate human interpretation [30]. Depending on the application, ML for biology may tend towards traditional ML approaches that have fewer variables and greater interpretability to aid mechanistic understanding or hypothesis generation [31]. There are also significant challenges in the application of ML to biological data sets because of their relatively limited size compared to other data domains [32]. In general, if there are not orders of magnitude more observations than predictors, complex deep learning models can underperform compared to more traditional ML algorithms [33].

We sought to further explore the connection between transcriptional immune responses and disease output in order to identify general patterns that predict final disease outcome using publicly available data. The initial research focused on the *A. thaliana-B. cinerea* pathosystem because the system is characterized by multigenic, small effect interactions, unlikely to be dominated by typical effector-NLR interactions [34,35]. Additionally, previous research has resulted in data sets large enough to employ ML [20]. Using a data-driven ML analysis pipeline, we tested if an ensemble of post-infection transcriptional responses could inform final disease outcome across a set of diverse pathosystems. For this case, ML is a powerful approach that does not require pre-selecting candidate genes or pathways, and it can capture complex, non-linear patterns across the whole transcriptome that collectively contribute to disease development, a complex multigenic trait. This approach can capture far more diverse, and likely biologically relevant, transcriptional patterns compared to traditional co-expression or differential expression analysis. Additionally, we employed a range of feature selection techniques, including those built from network theory and network geometry, to identify specific sets of genes providing accurate disease prediction across pathosystems. Thus, our approach is a novel application of ML and network science to plant-immunology, resulting in the discovery of genes not previously associated with plant immunity or pathogen response that may capture a general plant immune response.

## Results

### Machine learning can predict disease outcomes from transcriptomics of host-pathogen interactions

We sought to test the hypothesis that transcriptional patterns during the early stages of plant infection are predictive for final disease outcomes. In order to gain understanding from the modeling, and to account for data size, we mainly focused on ML algorithms with fewer variables and interpretable components. To identify gene sets that are robust predictors of plant disease, we utilized feature selection and cross validation on multiple pathosystems (Fig. 1A). A total of five ML algorithms were trained on *B. cinerea* infecting *A. thaliana* [20,36] (Fig. 1A). The data included 96 diverse *B. cinerea* isolates infecting *A. thaliana* ecotype Col-0, along with a salicylic acid (SA) receptor mutant, *npr1* that is defective in SA-induced defense [37–39], and a bioactive jasmonic acid (JA) receptor mutant, *coi1* that normally participates in E3 ubiquitin ligase mediated activation of JA defense responses [40,41]. Transcriptional responses during host infection were measured by RNA-seq at 16 hours post inoculation and lesion size for each interaction was measured at 72 hours post inoculation (Fig. 1A) [20]. Transcriptional responses (i.e., processed RNA-seq per gene) were used as predictors and final lesion size as the response to train two kernel-based ML algorithms, support vector machine (SVM) and linear SVM [42], two decision tree-based algorithms, random forest (RF) [43] and extreme gradient boost (XGB) [44], and a deep neural-network (DNN) [45] (see Methods for details). In order to reach our goal of developing predictive models that can be used across pathosystems, continuous lesion size numerical value (i.e., disease outcome) was converted to ten disease classes, as commonly used for plant disease ratings. This conversion used a data-driven approach to create balanced disease classes and to normalize the distribution of observations (see Methods for details). Disease class predictions using both the host and pathogen transcriptomes showed a clear association between the observed and predicted values for the hold-out test data (Fig. 1B and Fig. S1). To quantify the results more clearly, we calculated the difference between the observed disease class and the predicted disease class, which we termed class error. For example, a class error of zero means the observed and predicted classes were the same, while a class error of 3 means the observed disease class was three classes higher than predicted (i.e., the prediction underestimated disease outcome). Host genotype significantly influenced disease class prediction, and *npr1* infected plants produced more similar predictions compared to wild type infected *A. thaliana* than did *coi1* infected plants (Fig. 1C and Table S1). This result likely reflects the substantial shift in disease outcomes seen for *coi1* infection (Fig. S2), and is consistent with the previously reported importance of *A. thaliana COI1* mediated defense against *B. cinerea* infection [20]. The difference in prediction accuracy between host genotypes was consistent when both the plant and fungal transcriptomes were modeled together (i.e., dual), or if only one organism’s transcriptome data were used (Fig. 1C and Table S2). This suggests ML algorithms are sensitive to genetic perturbation, and the impact of the *coi1* mutation on transcriptional response and disease development were more difficult to predict. Interestingly, across all host genotypes and ML approaches, the average prediction accuracy was significantly higher when models were trained using transcriptional data from both the host and the pathogen compared to either alone, (72% compared to < 68% respectively) (Fig. 1D). For all measures of model performance, predictions using the combined host-pathogen transcriptome data were either statistically similar or better than predictions using only one organism’s transcriptome (Fig. S3). For analysis using host-pathogen transcriptomes, the XGBoost model performed the highest with a prediction accuracy of 72.3% (Fig. 1D). The RF model provided the highest accuracy (70.4%) when only considering the host transcriptome, while the results from the linear SVM provided the highest accuracy (70.5%) for pathogen alone analysis (Fig. 1D). Collectively, these results show that modeling the response of both species together provides the most accurate prediction of plant disease outcome. This reflects the importance of both actors in determining disease development and highlights the dynamic and complex nature of dual-species interactions such as between a host and pathogen.

**Figure 1.**
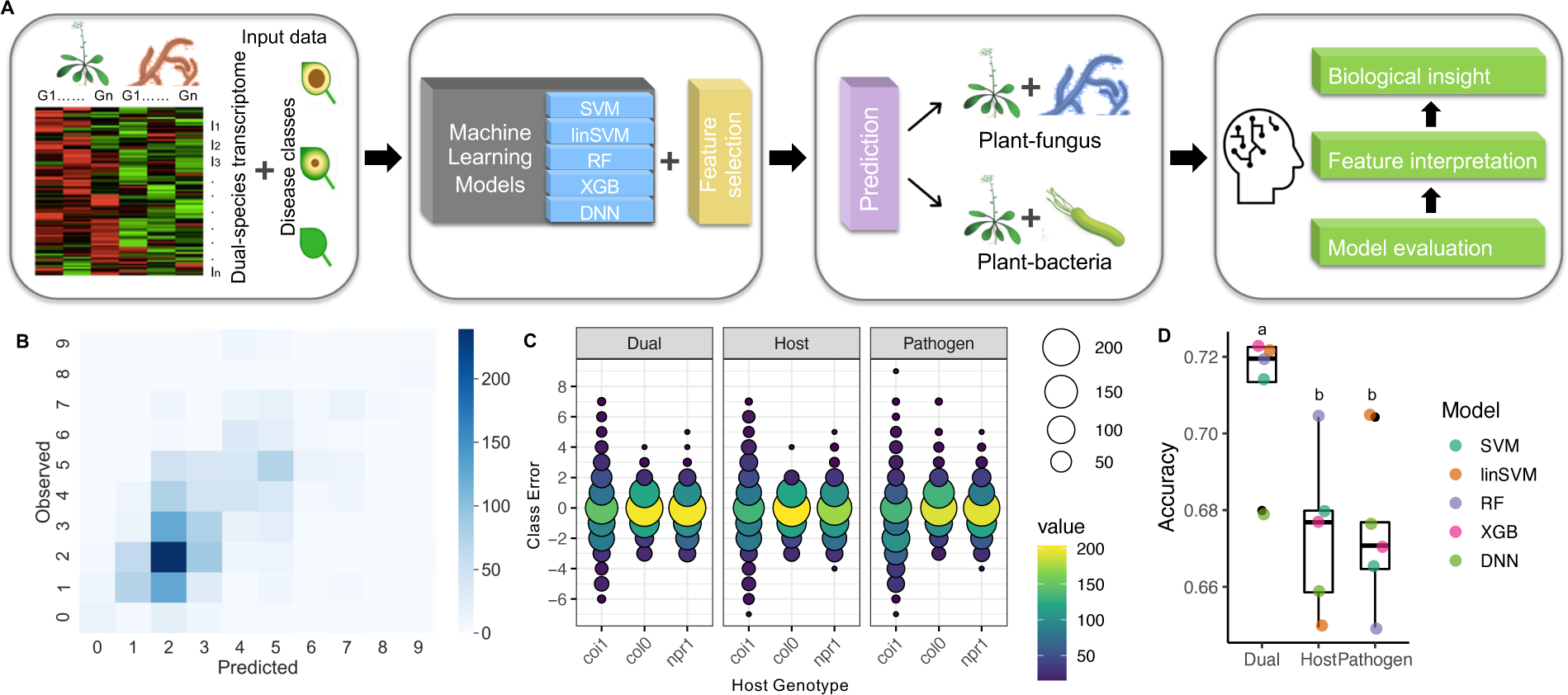
Machine learning can accurately predict later plant disease outcomes from earlier dual species transcriptome. (A) Schematic overview of interpretable machine learning strategies on plant immune elements in plant-pathogen pathosystems. (B) Heatmap of the observed and predicted disease classes based on the dual species whole transcriptome data. (C) Class errors calculated by predicted and measured plant disease classes on Arabidopsis-*Botrytis cinerea* transcriptome data [20]. (D) Accuracy of predicted plant disease class by five machine learning models either on the dual-species whole-transcriptome, or solely on plant host whole transcriptome, or solely on fungal pathogen whole transcriptome. Five machine learning models are support vector machine (SVM), linear SVM (linSVM), Random Forest (RF), XG Boost (XGB), and deep neural network (DNN). The letters above box blots indicate the significant levels (One-way ANOVA, p<0.05).

### Feature selection to identify general predictors of plant disease

We seek to identify a subset of transcripts that were both predictive of disease outcome, and that might also reflect meaningful biological mechanisms contributing to plant disease. Feature selection is a common practice in ML to help reduce the large predictors (*p*), small observation (*n*) problem associated with high-dimensional data [46,47] common in genotype-to-phenotype studies. A total of 12 feature selection approaches were employed using a range of techniques and feature set sizes to specify sets of transcripts used to train the ML algorithms and evaluate disease prediction. The feature selection approaches fall under the following six techniques-expert domain knowledge, statistical correlation, ML feature importance, co-expression network measures, co-expression network geometry, and a technique termed bipartite graph analysis (see Methods for details). For most feature selection techniques, the size of the feature set (e.g., how many genes to select) is not known. To experimentally determine this, we evaluated the impact of feature set size on model performance across seven feature set sizes ranging from 10 elements to the whole *A. thaliana* transcriptome (Fig. S4). The results indicate that both feature set size and model type impacted prediction accuracy (ANOVA, *P* < 2e-16, Table S3), and increasing the number of genes from 10 to 500 dramatically improved model performance (Fig. S4). We considered a number of set sizes for each feature selection method (Fig. S5), and the best feature set size for each feature selection approach as the inflection point for model performance across the tested feature set sizes (Fig. S6). For further evaluation, only a single set size for each feature selection technique was used, indicated as the number associated with the feature selection name. Evaluating disease prediction based on class error showed substantial variation in the agreement between observed and predicted disease classes, with both feature selection and ML model choice contributing to prediction performance (Fig. 2A and Table S4, S5). Predictions using the entire *A. thaliana* transcriptome resulted in the most zero class error predictions (i.e., no difference between predicted and observed), followed by the co-expression network degree set and the XGBoost feature importance set (Fig. 2A). Assessing the models for accuracy, eight of the feature selection sets performed statistically similar compared to predictions made using the entire *A. thaliana* transcriptome (Fig. 2B). This included an expert knowledge set based on previous characterization of *A. thaliana*-*B. cinerea* interaction [20,48], the two feature importance sets from RF and XGBoost, the two sets based on co-expression network measures, the two sets based on co-expression geometry, and the bipartite graph analysis (Fig. 2B). The goal to identify causal associations underlying feature sets must be assessed against random statistical associations in the data. To address this, we created random feature sets of 100 genes sampled from the *A. thaliana* genome, trained and evaluated model performance for the 5 ML algorithms, and repeated this process 100 times selecting new sets of random genes. Interestingly, we see that for four of the ML models, random gene feature sets can predict disease outcomes with approximately 60% to 68% accuracy (Fig. 2C). The SVM and RF had the highest average accuracies for the random gene feature sets at 65.7% and 65.4%, respectively (Fig. 2C). We interpret this result to show the power of ML models at identifying patterns in data, and in this case, biological meaning is not a prerequisite for predictive power, as has been noted previously [49]. Comparing the accuracy of the feature selection sets versus the results from many random sets of genes, many of the feature selection sets had a higher average accuracy than the random gene set, including Defense (67.0%), RF (67.4%), XGBoost (67.1%), Degree (66.2%), Betweenness (64.3%), NFD (67.9%), FDC (68.4%), and Bipartite (64.8%) (Fig. 2B). Assessing the ML models across all feature selections sets showed that the linear SVM and DNN consistently had lower prediction performance (Fig. 2D and E).

**Figure 2.**
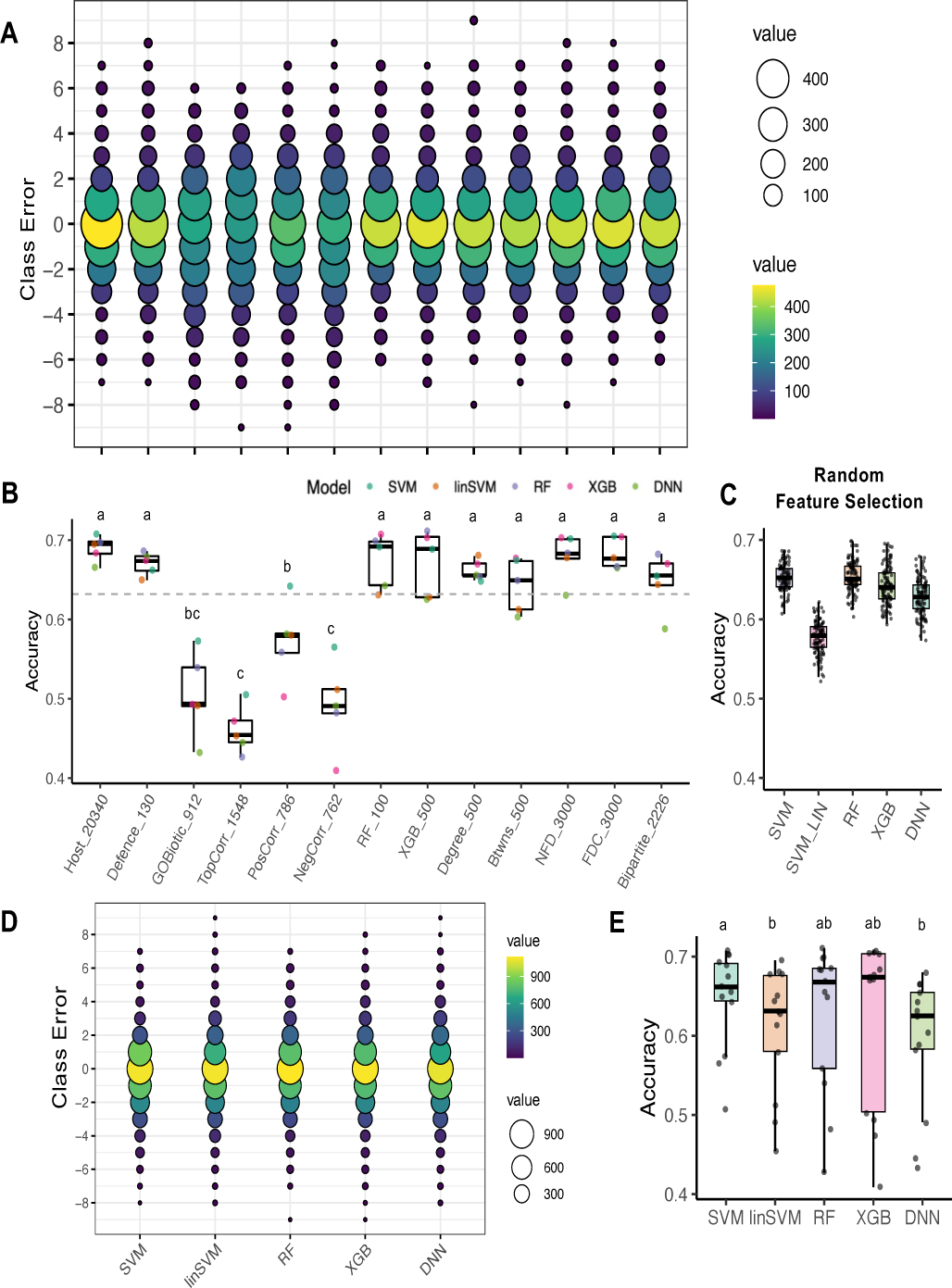
Evaluation of feature selection methods and ML models on *Arabidopsis* infected by *Botrytis*. (A) Class error of predicted and observed plant disease class. The resulting class error for each sample was counted and shown as a bubble plot, where the bubble size and color indicate the number of samples in that class. (B) Accuracy of plant disease class prediction by feature selection methods coupled for each of the five ML models tested. Support vector machine (SVM, green dot), linear SVM (linSVM, orange dot), Random Forest (RF, purple dot), XG Boost (XGB, pink dot), and deep neural network (DNN, light green dot). (C) Accuracy of the five ML models based on 100 randomly selected features over 100 iterations. Each dot indicates the accuracy from one set of 100 randomly selected genes for training and testing. (D) Class error of predicted and observed plant disease class as shown in A. Results are shown for all feature selection sets grouped by the five ML models. (E) Accuracy of predicted plant disease class of data shown in D. The letters above box blots indicate significance groupings (One-way ANOVA, p<0.05).

### Pretrained models can predict disease outcomes for new plant-fungal interaction

To further assess if specific feature selected genes can generally predict disease outcomes, we tested the models on new data not used for training. An independent data set of RNA-seq collected from *Arabidopsis* infected with *S. sclerotiorum* and non-inoculated control plants [50] was used to predict disease outcomes. The original disease classification was measured on a 1-6 scale, which we transformed to a 0-9 scale to be compatible with our pre-trained models (see Methods for details). The full analysis used the 65 trained models from the *A. thaliana*-*B. cinerea* interaction dataset, consisting of each of our five ML algorithms trained for each of the 12 feature selection sets plus the model trained on the entire *A. thaliana* transcriptome. Measuring class error across the models for each feature selection list showed that the bipartite feature selection list had the most zero class error predictions, followed by the GO Biotic response feature selection list (Fig. 3A). While many of the feature selection sets had a wide range of class errors, ten feature selection sets had the majority of predictions within one disease outcome class (i.e., between 1 and −1 class error; GOBiotic (63.3%), TopCorr (70.0%), PossCorr (63.3%), NegCorr (90.0%), XGB (73.3%), Degree (70.0%), Betweeness (56.7%), NDF (53.3%), FDC (56.6%), Bipartite (70.0%)) (Fig. 3A, B). Model assessment showed a wide variation of performance, driven by both the ML model and feature selection (Table S6). For example, the average accuracy across feature selection sets ranging from approximately 30% to 100% (Fig. 3B and Fig. S7). These results are also reflected in the 100 rounds of random feature selection assessment, showing a wide range of possible results across models, with the RF and SVM having the highest average accuracy (Fig. 3C). The two treatments, mock versus inoculated, had a significant impact on model performance, where uninoculated plants tended to have a positive class error while inoculated plants tended to have a negative class error, indicating an over and under prediction of disease development, respectively (Fig. 3D). The RF model had the highest accuracy across all feature selection sets, followed by predictions from the SVM algorithm (Fig. 3E). These results showed that ML models pre-trained on reduced feature selection gene sets from the *A. thaliana*-*B. cinerea* dataset, could correctly predict disease outcomes for an independent dataset of the same host infected with a different fungal pathogen, S. *sclerotiorum*. However, we note that the RF model had 100% prediction accuracy for 7 of 12 feature selection sets, and 5 of those 8 feature selection sets provided the highest accuracy disease predictions on the original *A. thaliana-B. cinerea* dataset (Fig. 2B and 3B). This suggests the ability to deliver highly accurate disease outcome predictions for independent pathosystems using ML, and that feature selection may allow a significant reduction in the number of predictor terms while maintaining model performance.

**Figure 3.**
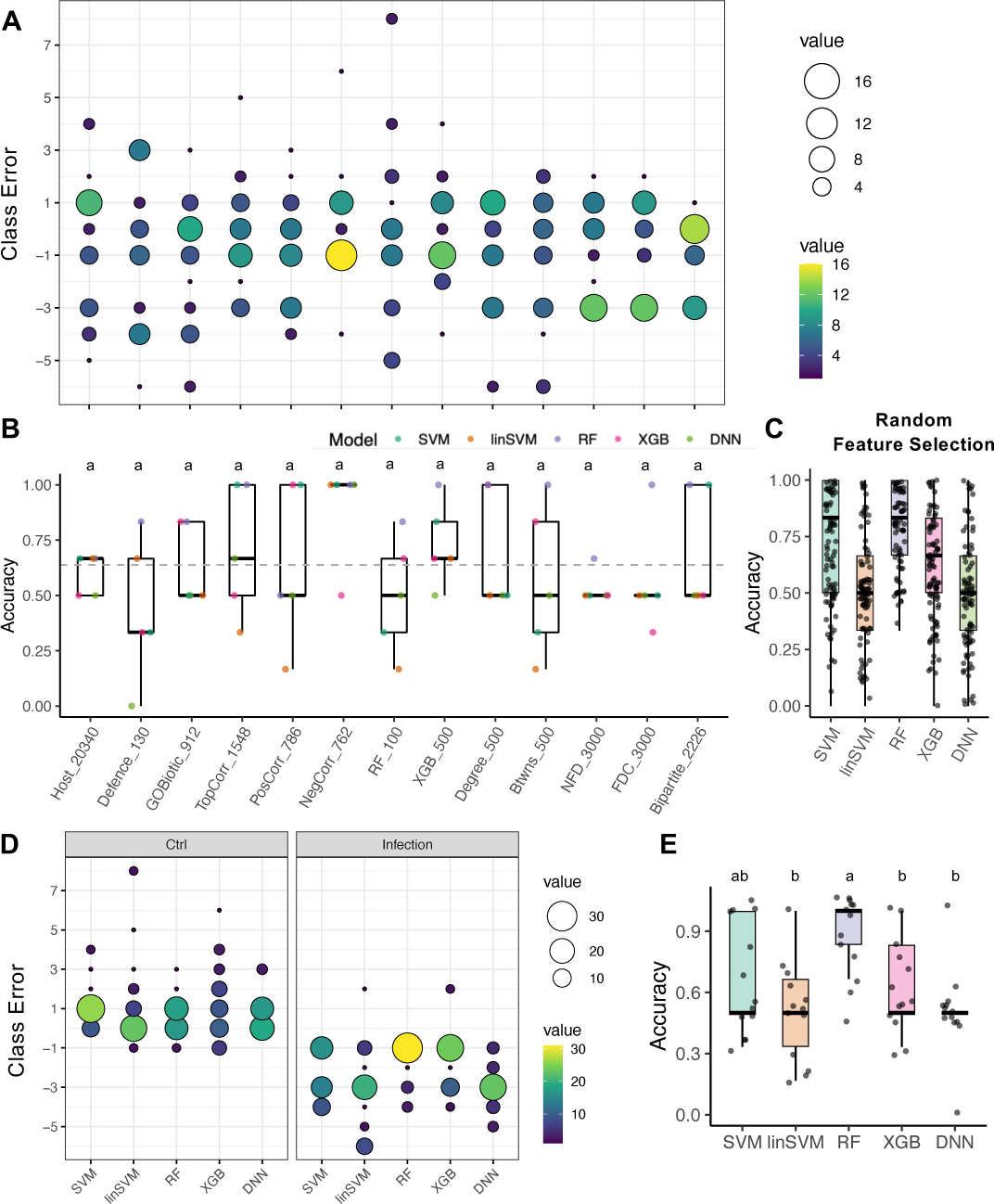
The trained machine learning models coupled with feature selection methods can accurately predict plant disease class on *Arabidopsis* infected by *S. sclerotiorum*. (A) Class error of predicted and observed plant disease class. The resulting class error for each sample was counted and shown as a bubble plot, where the bubble size and color indicate the number of samples in that class. (B) Accuracy of plant disease class prediction by feature selection methods coupled for each of the five ML models tested as previously described. (C) Accuracy of the five machine learning models based on 100 randomly selected features over 100 iterations as previously described. (D) Class errors calculated by predicted and measured plant disease classes on *Arabidopsis* transcriptome of control (Ctrl) uninoculated plants or infected (Infection) by *S. sclerotiorum* by five ML models. Results are shown for all feature selection sets grouped by the five ML models. (E) Accuracy of predicted plant disease class of data shown in D. The letters above box blots indicate significance groupings (One-way ANOVA, p<0.05).

### ML models can predict disease outcome for cross-kingdom pathogen attack

To rigorously investigate the modeling results, we extended our predictions to another independent data set, *Arabidopsis* infected with the bacterium *Pseudomonas syringae* [51]. The purpose of this analysis was two-fold. Firstly, changing host infection from an eukaryotic to prokaryotic pathogen broadly tests the robustness of the trained models and therefore their learned patterns. This addresses the issue of statistical pattern matching common in big data (i.e., overfitting), in which predictions are good for their original data, but poor for new datasets. If our models were overfit to the original dataset, then model performance for bacterial infection would be poor. Secondly, under the hypothesis that our approach identified general predictors of disease outcome, the models using feature selection sets would accurately predict disease outcomes for the independent data. The *A. thaliana* – *P. syringae* dataset comprised 27 different treatments made up of combinations of genetic differences in the host and pathogen, as well as immune priming (Fig. S8) [51]. The specific responses, such as immune priming and combinatorial immune mutants, would not be expected to produce transcriptional responses similar to the original training dataset, and therefore prediction results would likely not be accurate for such interactions. To test this, disease predictions were compared to the observed predictions and both host genotype and pathogen genotype impacted predictions (Table S7). The dataset was split for *P. syringae* expressing or not expressing AvrRpt2, and into *A. thaliana* wild-type Col-0, immune primed Col-0, and single, double or quadruple Col-0 immune mutants, and include all 5 ML models trained on the 12 different feature selection sets (Fig. 4A). The interaction between wild type host and pathogen produced substantially more zero class error predictions (32.8%), and predictions within one error class (46.11%), consistent with our expectations for better predictive performance (Fig. 4A). Compared to wild-type Col-0 infection with virulent *P. syringae*, disease severity was overestimated for the immune primed infection (i.e., flg22 pre-exposure), and underestimated on hosts with compromised immunity (Fig. 4A). Clearly, immune priming and genetic perturbation of host immunity negatively impacted predictive performance. The impact of immune priming and combinatorial genetic perturbations was also seen using accuracy assessment of class error, showing that non-wild type interactions negatively impacted model performance (Fig. 4B). Prediction accuracy was also lower when *A. thaliana* was infected by an avirulent *P. syringae* strain expressing *AvrRpt2* (Fig. 4C). Collectively, these results show that disease predictions for wild type compatible host-pathogen interactions were more accurate than predicting incompatible or host immune compromised outcomes. This suggests that genetic perturbations and NLR based immunity caused a transcriptional profile that was too dissimilar from the original training data for accurate prediction. Therefore, only data for virulent *P. syringae* infecting wild-type Col-0 (13 observations) were used for further analysis.

**Figure 4.**
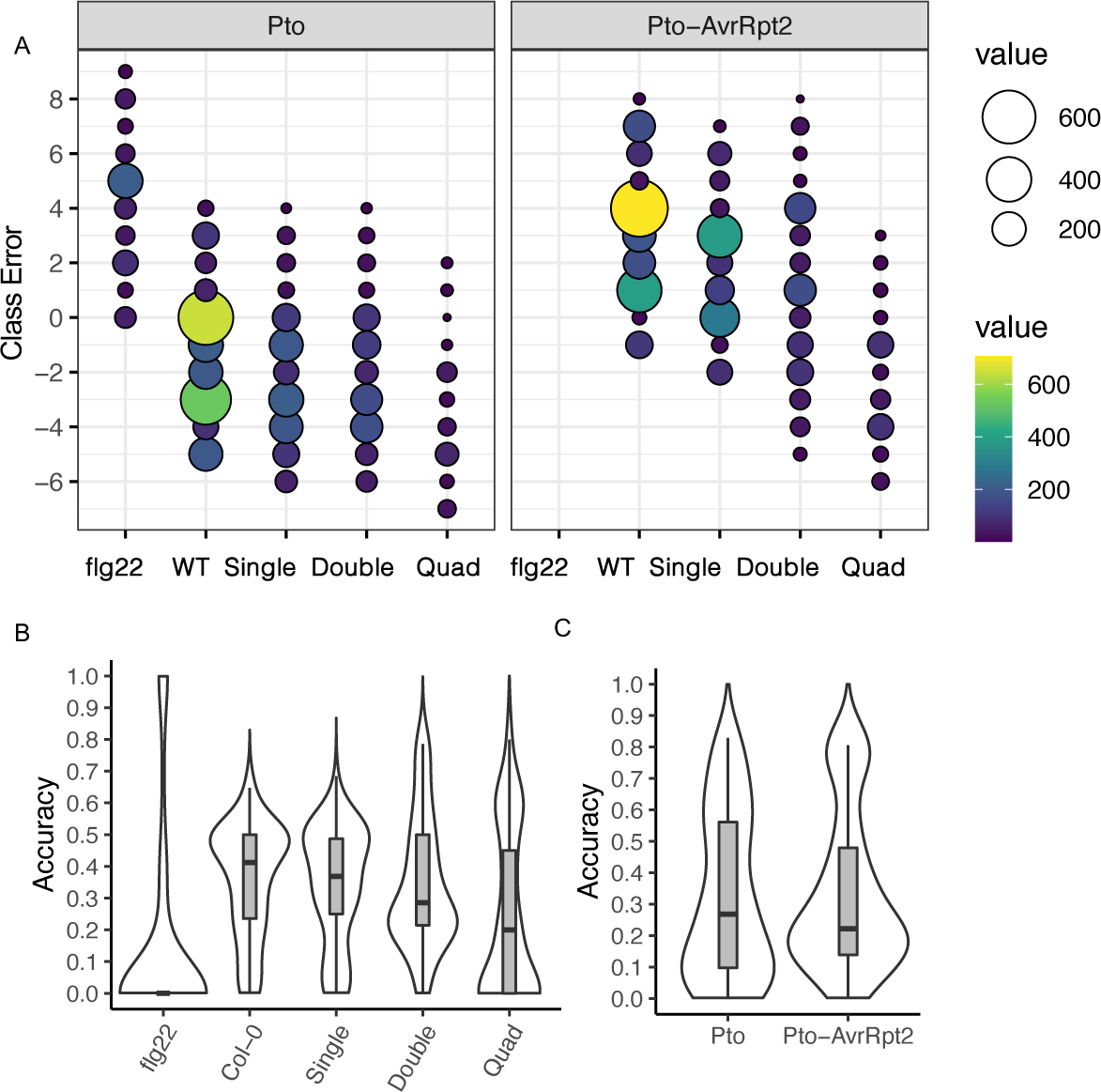
Genetic perturbations for immunity and virulence hamper generalization for bacteria disease prediction. (A) Class error of predicted and observed plant disease shown as a bubble plot. Individual host treatments were grouped by immune priming (flg22), wild type Col-0 (WT), or single, double, or quadruple mutation in Col-0. Interactions with Pto *P. syringae* shown in left plot, and infection with Pto *P. syringae* expressing the avirulence gene *AvtRpt2* shown on right. Infection by Pto-AvtRpt2 did not use immune priming as a treatment. The results were combined from predictions using all 5 ML models and feature selection sets previously described. (B) Accuracy of predicted plant disease class grouped by host treatment. (C) Accuracy of predicted plant disease class grouped by pathogen treatment.

To understand how our *A. thaliana-B. cinerea* trained models with feature selection perform on *A. thaliana-P. syringae* wild-type interaction data, disease predictions were generated. Assessment through class error showed variable results across feature selection set trained models (Fig. 5A). The feature set based on network analysis node betweenness gave the most zero class errors predictions (Fig. 5A). Predictions using feature selection sets Defence (53.3%), RF (68.1%), and Betweenness (74.8%), provided at least half of their respective predicted disease classes within one class error of the observed value. Quantifying prediction accuracy also indicated variable performance across the ML models (Fig. 5B). These results reflect the complexity of the prediction problem, which uses host transcriptome responses of only a small subset of genes from early infection of a fungal pathogen to predict disease outcomes for bacterial infection. Nonetheless, 6 of the 12 feature selection sets produced an average accuracy greater than the average random feature set prediction accuracy (36.6%) (Fig. 5B, C): Defence (48.9%), PosCorr (42.2%), NegCorr (40.0%), RF (74.4%), Betweenness (72.2%), and Bipartite (46.7%). Also, 4 of the 8 feature selection sets that performed above the random feature sets for *A. thaliana-P. syringae* were the same that provided the highest accuracy predictions from the original *A. thaliana*-*B. cinerea* training (Fig. 2B and 4B). This is an indicator of robust performance for these feature selection sets. Looking at individual model performance, the distribution of predicted class errors and prediction accuracy shows that some ML algorithms performed better (Fig. 5D and 5E). When analyzed across all feature selection sets, the RF model had the highest prediction performance, average accuracy of 65.4%±28.0% (mean±SD), followed by XGBoost, average accuracy 58.5%±34.4% (mean±SD) (Fig. 5D, 5E). Overall, the change from a eukaryotic to prokaryotic pathogen did not preclude accurate disease prediction, suggesting that a common transcriptional response could be modeled that predicted disease outcomes across a range of interactions.

**Figure 5.**
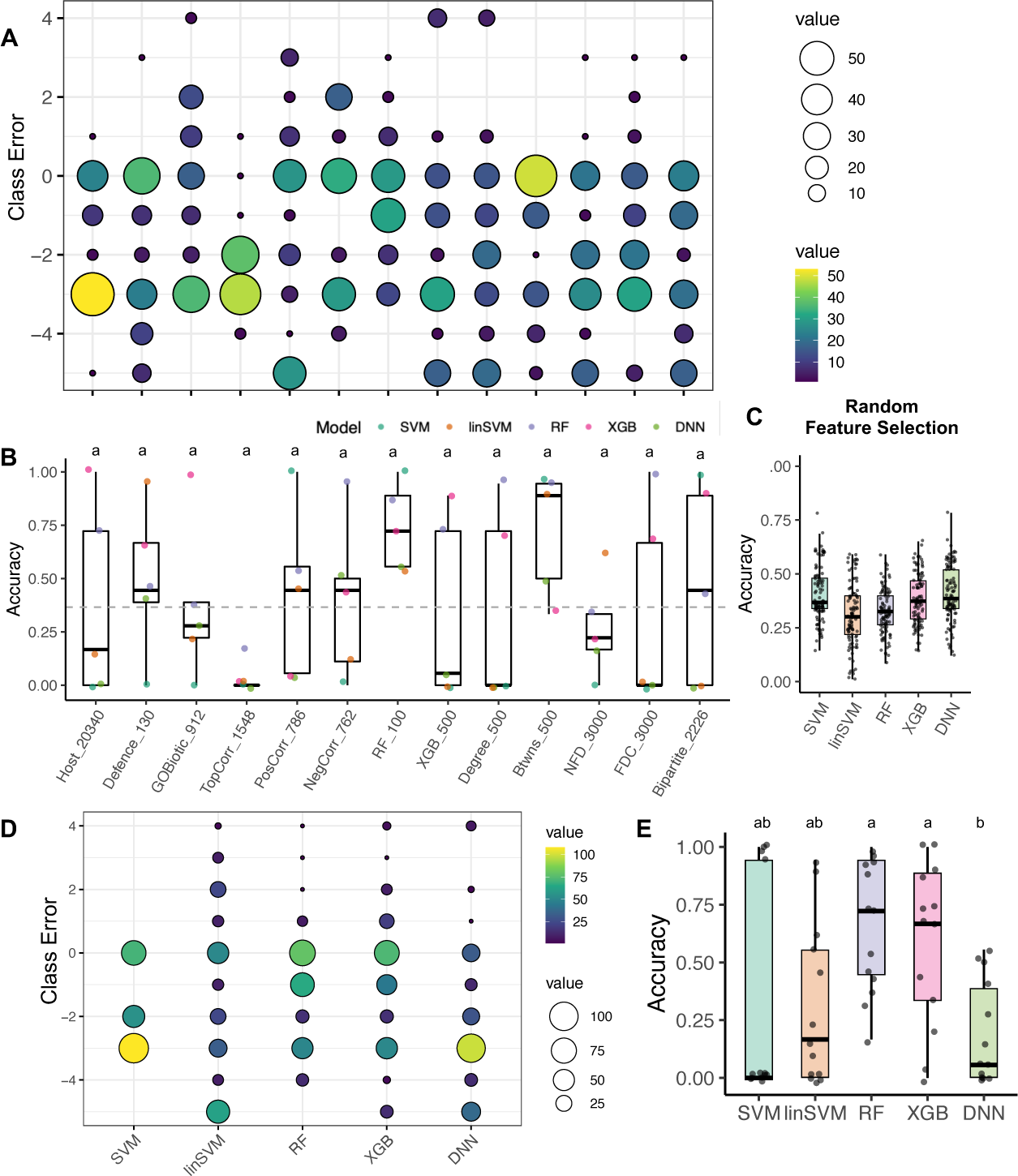
Trained ML models can predict disease outcomes for new pathosystems. (A) Class error of predicted and observed plant disease class. The resulting class error for each sample was counted and shown as a bubble plot, where the bubble size and color indicate the number of samples in that class. (B) Accuracy of plant disease class prediction by feature selection methods coupled for each of the five ML models tested as previously described. (C) Accuracy of the five machine learning models based on 100 randomly selected features over 100 iterations as previously described. (D) Class errors calculated by predicted versus observed disease classes for all observations by the five ML models. Results are shown for all feature selection sets grouped by the five ML models. (E) Accuracy of predicted plant disease class for data shown in D. The letters above box blots indicate significance groupings (One-way ANOVA, p<0.05).

### A novel set of transcripts accurately predicts plant disease across a range of pathosystems

We aimed to further understand the feature selection sets and the underlying genes. To integrate the analysis across all three pathosystems, we re-focused our analysis on results from only the RF and XGBoost algorithms that proved most accurate for *P. syringae* predictions (Fig. 5E). Disease predictions across the three pathosystems were re-assessed for the RF and XGB trained models for each feature selection sets (Fig. 6A). The average accuracy for all feature selection sets is higher than the random feature selection set (Fig. 6A) (random feature selection accuracies for RF (32.4%) and XGB (38.4%)). Remarkably, three feature selection sets, XGB (79.1%), Degree (80.7%), and Betweenness (77.4%), have an average prediction accuracy of over 75% across all three pathosystems (Fig. 6A). We did not find KEGG pathway enrichment for any of the feature selection gene sets. The transcriptional response of each gene set per host infection indicated that these genes are not necessarily the most highly expressed genes during infection (Fig. S9A), nor are they significantly more induced as a group during infection compared to non-infected (Fig. S9B, C). These results are interesting because most transcriptional analysis of the host immune response involves identification of highly expressed genes or genes differentially expressed between healthy and infected plants. Comparing the individual gene composition for each feature selection set indicates that they are largely non-overlapping sets (Fig. 6B), indicating that the different methods identified different genes. Two gene sets rom previously published literature were used as controls to understand and validate the results. One control gene set contained 2524 genes previously identified through genome-wide association study (GWAS) of *A. thaliana* genotypes infected by *B. cinerea* [48]. A C*hi-square* test of independence showed that the GWAS gene set significantly overlapped the feature selection set identified through the XGBoost model, bipartite graph analysis, and the positive correlation analysis (Fig. 6C, and Table S8). We also compared the feature selection sets to 970 genes identified as common early transcriptionally responsive genes in *A. thaliana* to diverse immunogenic elicitors [18], referred to, here, as the Bjornson set. This set of genes was remarkably overrepresented in nearly all of our identified feature selection sets (10 of 12 sets), except that identified from XGBoost and the bipartite graph analysis (Fig. 6B, and Table S9). These validation experiments indicate that our identified feature selection gene sets are significantly enriched in genes previously identified for resistance to *Botrytis* or PRR-induced immune signaling. These results support the hypothesis that the feature selection gene sets represent collections of genes broadly predictive of Arabidopsis disease development, and that a portion of the genes may have biologically relevant roles in disease development.

**Figure 6.**
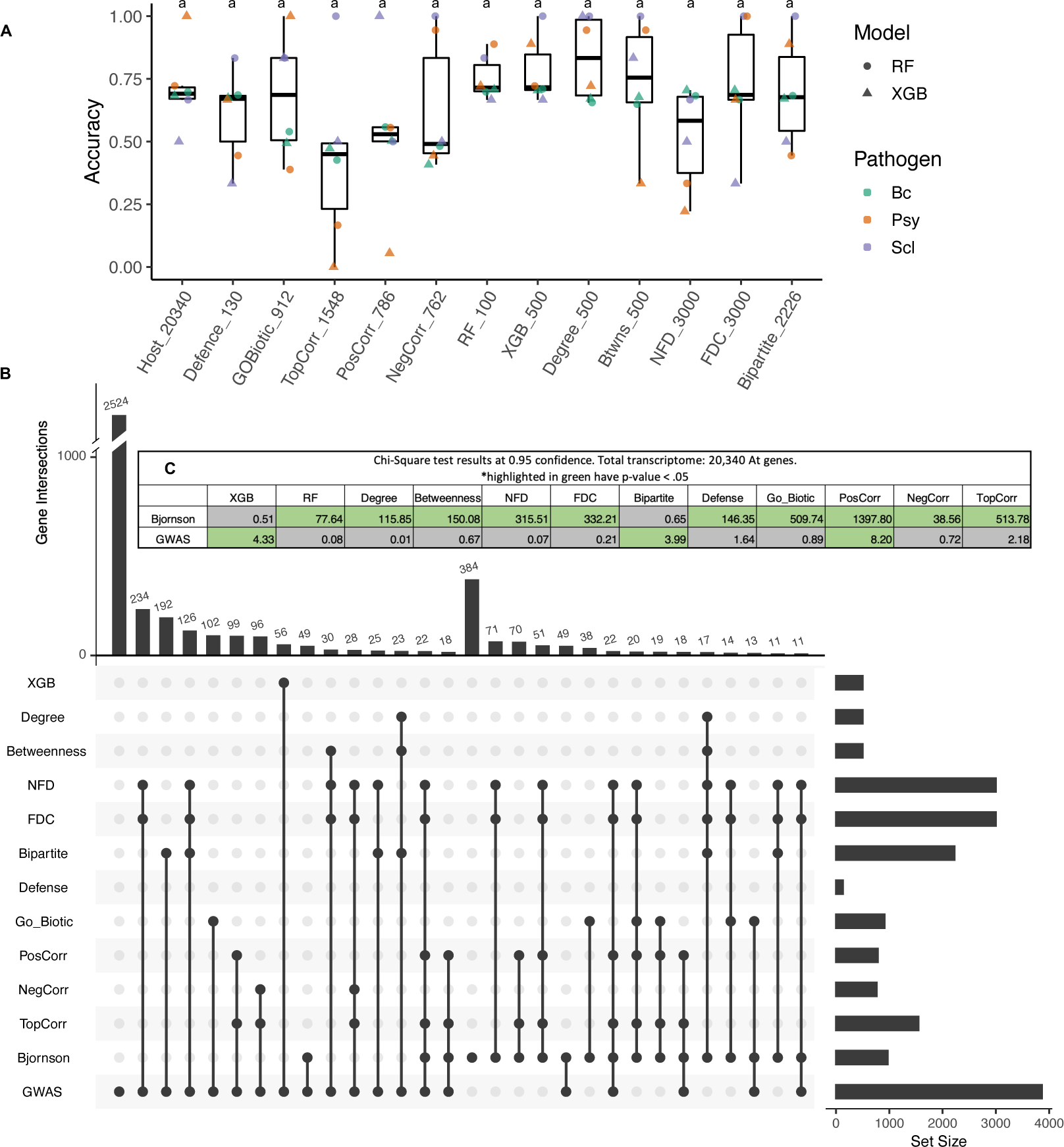
Uncovering general predictors of the plant immune response. (A). Prediction accuracy for each indicated gene set from the random forest (RF, circle) and extreme gradient boost (XGB, triangle) models across infection data sets *B. cinerea* (Bc, green), *P. syringae* (Psy, orange), and *S. sclerotiorum* (Scl, purple). (B) UpSetplot of *A. thaliana* gene features based on different feature selection lists. This shows the top 15 set intersections between the different feature selection lists and the Bjornson [18] and GWAS [48] gene lists. (C) Table of chi-square test of independence at 0.95 confidence for the different feature selection lists versus the Bjornson and GWAS gene lists.

## Discussion

Plants possess two distinct classes of immune receptors residing at the plasma membrane or in the cytosol that function to detect and counteract microbial reproduction in the host [52]. Evidence indicates that the plant immune system can be conceptually thought of as an integrated functional unit, with cross-talk between immune receptor classes and disease outputs providing synergy, redundancy, and specificity [13,53,15]. Despite our growing knowledge of the plant immune system, there are few tools or approaches that can predict a plants general immune performance. Current research and improvement approaches rely heavily on specific single-gene interactions, which have proved effective, but are hampered by their intensive time requirements and lack of generalization. Additionally, such genetic resistance has proven relatively short lived when deployed at scale [54], although considerable effort is being put into stacking immune receptors to increase efficacy in the field [55–57]. As plant sciences and crop improvement move to engineered systems approaches [58–60], greater understanding and predictive power for processes such as plant immunity, abiotic stress tolerance, growth, and nutrient utilization are needed. Systems approaches leveraging increasingly common big data sets, analyzed by interdisciplinary teams, can help provide the knowledge and framework to achieve these goals.

Here, we addressed general plant immunity by looking for early transcriptional patterns that are predictive of disease outcomes across a range of pathogen attack. Our hypothesis was that plants have a common early signaling response to a range of immunogenic signals that lead to different degrees of defense output. We tested this hypothesis by training models on RNA-seq data from one pathosystem, and independently testing the trained models on new input data from different pathosystems. Such a model should only be predictive of disease outcome in the new system if the model learned general patterns of early RNA-seq response indicative of final disease outcome. We further refined our approach, and addressed the underlying biology, by identifying ten subsets of host genes using feature selection techniques in order to train the ML models with fewer parameters. A significant finding from this research is that ML models trained using only a fraction of the total host transcriptome, from 0.5% to 15% of total genes, were able to accurately predict disease outcomes across fungal and bacterial pathosystems, including both necrotrophic and biotrophic pathogens. This is important because many ML models do not perform well on new data sets, showing a lack of generalization. We interpret this high predictive performance across diverse data sets to reflect a general disease-related transcriptional response that was captured by the models. It is interesting that different feature selection gene sets, that have largely non-overlapping membership, can independently provide high-predictive power across diverse pathosystems. This result is consistent with the idea of immune response canalization, and that immune signaling is a robust network with many paths to a similar response [13,20].

There are many novel aspects to our ML interrogation of the plant immune response. Our approach overcomes general limitations for the most common analysis pipelines, such as differential gene expression (DGE), cluster or module analysis, and network construction utilizing simple co-expression (i.e., correlation). Such approaches fail to capture non-linear relationships, have limited predictive power, and have limited generalizability. For instance, commonly used DGE simply reports transcript levels that are higher or lower between conditions, but the approach fails to integrate patterns or relationships between transcripts which collectively account for a response of interest. Gene co-expression networks integrate relationships between transcripts, but rely on simple linear correlations between transcripts, and assume that transcripts with similar patterns are functional related. In contrast, ML models can identify more diverse transcriptional patterns that collectively correspond to phenotype development. The ML models employed here capture non-linear dynamics, are not limited to identifying transcriptional outliers, and inherently integrate multigenic patterns. Another contribution of this research is demonstrating how to link biological questions to the deployment of ML to enhance our understanding and interpretation. A concern for ML in biology is the identification of biologically irrelevant patterns. This can occur during model training, where patterns particular to a given data set are learned (i.e., overfit). We reasoned that any ML model that was overfit on a particular dataset would have limited predictive power on new independent data. Therefore, we were able to validate and identify which feature selection gene sets are the most likely to represent biologically meaningful patterns. However, we note that this does not mean the identified transcripts and patterns are directly related to plant immunity. They may function in totally unrelated pathways, but have feedback or secondary interactions with the immune response, and powerful ML algorithms can learn such patterns that are predictive of the final outcome. Such a situation is a problem if the research goal is to leverage information from the model to change the system. For example, gene products and patterns of transcription that are predictive of low disease development could be used as breeding targets for crop improvement, but the approach will likely only be successful if the pattern is directly related to cause or mechanism. Future tools and research approaches are needed to further develop robust design-build-test-learn pipelines for routine use.

There is increasing interest in predictive biology and engineered systems to benefit society. We provide here an example of mining publicly available data to understand general components of plant disease development. This approach is general and scalable, allowing for the interrogation of other data sets and integration of new data as it becomes available. It will be of great interest to go from predicting disease outcomes from RNA-seq to predicting general disease outcome from DNA sequence. Resulted presented here provide new understanding of complex plant-microbe interactions and offer a framework for future research.

## Methods

### Machine learning tasks

To understand the relationships between transcriptional response and disease outcome, we used the state-of-the-art supervised machine learning algorithms to train models that can accurately predict plant disease severity from gene expression data. The classification task is to predict the disease severity (classes) based on the gene expression profile of dual or sole species. We used plant disease phenotypic data, fungal colonized lesion area in plant-fungal pathosystems or bacteria growth in plant-bacteria pathosystems, as labeled multi-class data. The transcriptomic data derived from dual species or solely from plant host or pathogen serve as input data. To identify the crucial gene set that control disease development, we performed different feature selection methods on the training plant gene expression datasets and applied the selected genes on different trained models to evaluate performance improvement. The multi-step workflow starts from plant disease phenotypic data and RNA-seq read counts data, followed by ML and feature selection methods, and outputs disease predictions (Fig. 1A).

### Disease phenotypic and transcriptomic data acquisition

We used the previously published plant disease phenome and transcriptome data involving 1,164 *Arabidopsis* and *Botrytis* interaction pathosystems [20,36]. In brief, the *Arabidopsis-Botrytis* dataset include four replicates of 97 Botrytis isolates infecting three *Arabidopsis* (Col-0) genotypes-wild type with complete immunity, and two immune compromised lines, *npr1* and *coi1,* compromised in SA and JA mediated defense response respectively. The plant disease phenotypic data, fungal colonized leaf area, were measured at 72 hour-post inoculation. The transcriptomic data were collected at 16 hour-post inoculation and raw RNA-seq data are available from the National Center for Biotechnology Information (NCBI) under BioProject no. PRJNA473829 and accession no. SRP149815.

The validation set of plant-fungal pathogen interaction was derived from *Arabidopsis* and *Sclerotinia sclerotiorum* interactome [50]. The disease phenotypic data were collected at 24 hour-post inoculation, and tissues at the edge of developed necrotic lesions were collected for RNA extraction. Data were accessed through NCBI Gene Expression Omnibus (GEO) accession GSE106811. The validation set of plant-bacteria pathogen interaction was derived from *Arabidopsis* and *Pseudomonas syringae* interactome [51]. The disease phenotypic data were measured as the bacteria growth at 48 hour-post inoculation. The transcriptomic data were collected at 6 hours-post inoculation and accessed through are available through NCBI GEO GSE103442. We note, there was a problem with miss-labeled metadata samples for the pre-treatment samples. Only samples pre-treated with flg22 were retained for initial analysis, the rest of the samples receiving pre-treatment with chitin or SA were dropped due to ambiguous labeling in the metadata file on SRA.

### Data processing

To perform the multi classification learning task, we divided the *Botrytis* infecting *Arabidopsis* training disease phenotypic data into 10 classes based on the disease severity to generate multiclass labels. We consider the largest lesion from the training dataset corresponding to the highest disease class (Class 9). A lesion size of 0 is mapped to the lowest disease class (Class 0). Then, we divided the lesion data into 10 equal lesion bin sizes and labeled each training sample accordingly. To obtain comparable disease class label for the plant-fungal pathogen test data, the labels for healthy samples and infected samples were transformed as disease class 1 and 6 based on the distribution patterns of lesion area measured from *Arabidopsis*-*Sclerotinia* pathosystems and *Arabidopsis*-*Botrytis* pathosystems (Fig. S2). Similarly, disease class labels were assigned based on the comparison of distributions of lesion area from *Arabidopsis*-*Botrytis* pathosystems and bacteria growth measured from *Arabidopsis*-*Pseudomonas* pathosystems (Fig. S2 and S8).

To avoid the bias caused by the multiple mapped reads to both plant host and pathogen reference genomes, we used dual-species reference genome to map the RNA-seq reads derived from the pathosystems [61,62]. Briefly, we first generated a dual species reference genome by concatenating the *Arabidopsis* TAIR10 reference genome [63] with either of the three pathogen reference genomes, *B. cinerea* (strain B05.10, build ASM83294v1) [64], *S. sclerotiorum* (strain 1980 UF-70, build ASM185786v1), or *P. syringae* (strain *Pseudomonas syringae* pv. *tomato* DC3000, build GCF_000007805.1) [65] respectively. Raw RNA-Seq reads were mapped to the corresponding combined dual species reference genome by STAR [66]. Raw gene counts were normalized as transcripts per million (TPM) [67]. For the *Arabidopsis-Botrytis* data set, individual sequencing libraries were removed if they had < 30% uniquely mapped reads. Annotated genes in from *Arabidopsis* were removed from the count table if the gene had read counts < 200 from across all sample libraries. After quality control, there are 1,102 samples and 20,340 expressed *Arabidopsis* genes 8,761 expressed *Botrytis* genes included in the *Arabidopsis*-*Botrytis* training dataset. The same criteria were used to assess the two validation data sets (*A. thaliana – S.sclerotiorum* and *A. thaliana - P. syringae*), and both sets passed quality control. To be consistent, only count data for the 20,340 *Arabidopsis* genes used in training were retrieved from the validation data.

The individual ML models were trained on the gene expression values of the Arabidopsis-Botrytis dataset. Data were split into 70% training and 30% test sets. Since the range of gene expression values vary significantly per gene, feature scaling is needed to ensure that the contribution of each feature is not biased towards larger numerical values. In addition, scaling the features ensure faster gradient descent convergence for some of the machine learning models. Here, we applied min-max standardization [68]. Additionally, the Arabidopsis-Botrytis dataset is highly imbalanced with respect to the lesion size/disease class (Fig. S2). Lower disease classes (class 0-3) are more represented in the data than higher disease classes (class 6-9). To mitigate this data imbalance, we utilized SMOTE (synthetic minority oversampling technique) to oversample the minority data classes to have an even representation of the classes during training [69]. This technique increases the minority class examples by synthesizing new data stochastically from the existing training class sample space. Note that it is important that the training/validation/test splits should be done before any preprocessing steps (e.g., data standardization) to avoid “data peeking” which can over-inflate test prediction performance. For example, data standardization of the validation data should be done based on the aggregate statistics of the training data only. If the average or data statistics is based on the entire dataset (including validation and test sets), in effect we have considered information from the test data.

### Supervised model training

According to the multi-classification task and the data characteristics in this study, we selected supervised machine learning strategy to build the learning models that can learn from the transcriptome input derived from dual species or sole species and can accurately predict the correct phenotypic disease class outcome in a plant host and pathogen interaction pathosystems. A total of five popular ML algorithms for supervised learning were tested, including two support vector machine-based algorithms, two decision tree-based algorithms, and a deep neural network algorithm. Hyper-parameters tuning was performed for each model. Support vector machine is a supervised learning algorithm that can efficiently perform both linear and non-linear classifications on data for robust prediction. It can construct a hyperplane or set of hyperplanes in a high-dimensional feature space to achieve the largest distance to the nearest training-data point of any class. Although support vector machines are naturally used for binary class tasks, the algorithms can be developed to apply on multi-classification tasks by reducing the multi-class task to several binary problems [42]. Linear support vector machine is a linear classifier that is used for linearly separable data into classes by using several straight lines. We used support vector machine and linear support vector machine, SVC(), from scikit-learn package [70,71]. The parameters for support vector machine and linear support vector machine are based on the default setting.

Decision tree-based algorithms generally have high performance on small-to-medium structured data, and are a popular and robust approach for various machine learning tasks because it is invariant of feature scaling and transformation, and independence of irrelevant features ensure constructing inspectable models [72]. Decision trees makes decisions using a graph to represent of all possible solutions queried by certain conditions. Random forests, also called random decision forests, are decision-tree-based ensemble algorithms that perform classification tasks using a bagging strategy to build a multitude of decision trees (forest) where only a subset of features is randomly selected to build a forest or decisions are collected from some trees [43]. Therefore, the accuracy of such ensembled decisions by random forests is generally higher than models built on single random decision tree. However, random forests show biases in data including categorical variables with high variation levels and/or correlated variables, which are common cases in transcriptomic data. We used random forests (RandomForestClassifier) for learning modeling training from sklearn python package with parameters n_estimators=100, max_depth=30, random_state=0. Furthermore, we used random forests for ranking the importance of features for feature engineering.

Extreme gradient boosting (XGBoost) is another decision-tree-based ensemble learning algorithm that uses an optimized gradient boosting algorithm for classification, regression, and ranking tasks. Gradient boosting builds sequential models by using gradient descent algorithm to minimize the errors from previous models while increase the influences of high-performance models [44]. Such strategy can solve the drawbacks of random forest-based models learning from data with high varied level of categorical variables and/or correlated variables. Based on gradient boosting framework, XGBoost is further optimized to avoid overfitting and/or bias through several ways, including parallel processing, tree-pruning, handing missing values and regularization (12). Furthermore, it dramatically improves the computing power for boosted tree algorithms. We used XGBoost (*XGBClassifier*) for modeling training from XGBoost python package with default parameters. The ranked features generated from the XGBoost model are used for feature selection. We also included an artificial neural network-based model among our supervised learning algorithms for comparison. We used *Keras* package to build the modular neural network available from *TensorFlow* package [73] to build a deep neural network model with a substantial credit assignment path of depth 2 (with batch normalization, 512 dense hidden layer with ReLU activation function, dropout = 0.5, dense 10-class output layer with softmax activation function). The cross-entropy loss and the Adam optimizer were used with a learning rate of 0.001. The trained neural network model could directly transform the raw whole transcriptome input into a more abstract and composite intermediate feature representation and extract such higher-level intermediate features to predict disease severity.

### Feature selection

To better understand genes contributing to plant disease development, we performed feature selection on expressed genes from the whole *A. thaliana* transcriptome. Feature selection methods were assessed by using the feature selection gene sets in each trained model to observe performance improvement or decay. The full set of feature selected genes by the different methods can be found in Table S4. The plant whole transcriptome data served as a baseline control using the random forest model with a prediction accuracy of 70.4%. We randomly selected 100 plant transcripts as negative control with the highest prediction accuracy of 65.7% by support vector machine model. We explored domain knowledge-based methods to obtain the plant genes associated with pathogen defense. The list of 130 defense genes is based on previously published literature characterized as having defense function in the *Arabidopsis-Botrytis* pathosystem. The term ‘response to biotic stimulus’ from Gene Ontology was used to identified 1,182 *Arabidopsis* genes [74], of which, 912 had detectable expression in the *Arabidopsis-Botrytis* transcriptome. Previously published *Arabidopsis* genes that were either positively or negatively correlated with plant disease lesion areas were selected as feature selection sets PosCorr_786 and NegCorr_762 [20,36]. The RF_100 and XGB_500 feature selection sets were generated from the plant gene set based on the ranked importance during model training.

To obtain the gene set based on the gene co-expression network topology, we first constructed the plant gene co-expression network using the *Arabidopsis* transcriptome data. We then calculated the network structure parameters, including node degree (Degree_500) measuring the number of connections between a node (gene) and its neighbors, betweenness centrality (Btwns_500) measuring the number of shortest paths between all pairs of nodes in the network which pass through the focal node. Apart from the classical network metrics, we used for node ranking and feature selection the fractal dimension centrality (FDC_3000) and node fractal centrality (NFD_3000), which captures the topological features and the degree of complexity and heterogeneity in generating rules of complex networks [75]. Additionally, we also used the bipartite graph-based feature selection method to extract the *Arabidopsis* genes based on their co-expression relationships during *B. cinerea* infection. First, the bipartite network is created from the plant-pathogen co-expression dataset. Then, a unipartite network projection on the *Arabidopsis* genes is extracted from the bipartite network based on the frequency of shared connection to the *Botrytis* gene set [76]. We assessed feature selection membership using an UpSet plot from package (()). Chi-square test of independence were used to evaluate dependence between the feature selection sets and two gene sets previously implicated in GWAS host response [48], or general response to immunogenic peptides [18].

### Evaluation

We measure the model performance based on the classification errors. Since a +1/-1 classification error is within the tolerable error range, we used the 1-class error True Positive (TP_1_) to define the accuracy of the predicted disease classes. We used a confusion matrix approach to evaluate the performance of the multi-classification task of machine learning models and feature selection methods. Each prediction falls into one of the four cases: true positive (TP_1_), false positive (FP), true negative (TN), and false negative (FN). To describe distribution of predictions across each of these categories, we calculated a variety of performance matrix, including Accuracy, Precision, Recall, F1_Score, and mean squared error (MSE).

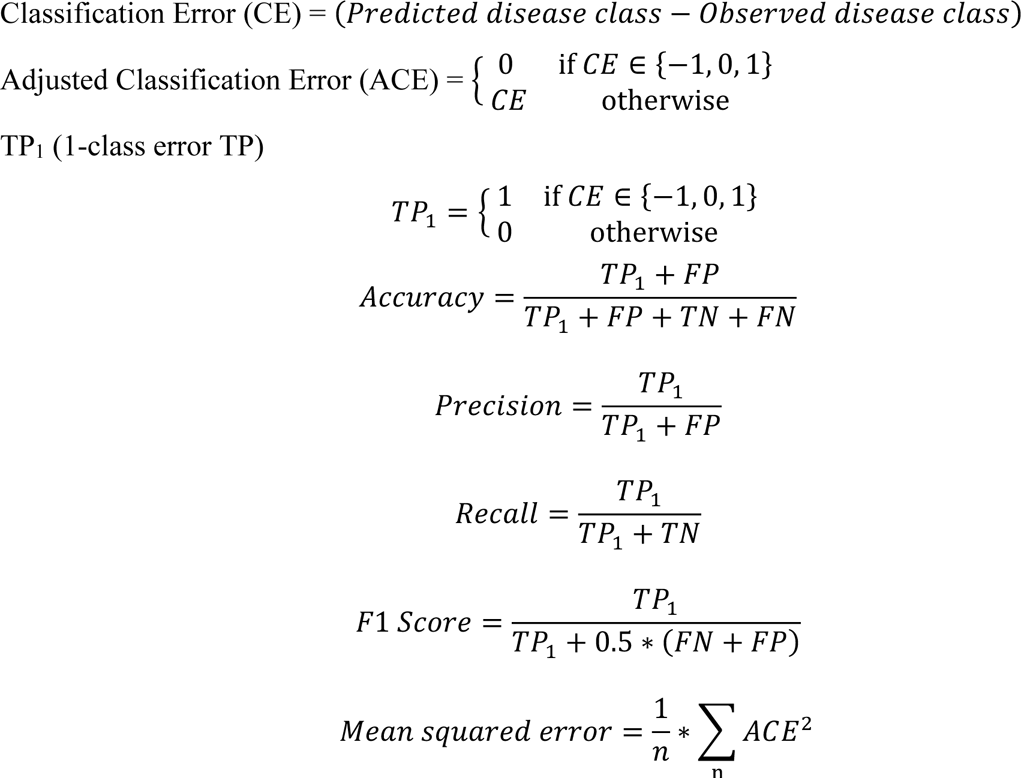

## Data availability

All datasets used in this study are publicly available. The *A. thaliana* and *B. cinerea* transcriptome data are available through NCBI Bioproject PRJNA473829 [20,36]. The *A. thaliana* and *S. sclerotiorum* transcriptome data are available through NCBI GEO accession GSE106811 [50]. The *A. thaliana* and *P. syringae* transcriptome data are available through NCBI GEO GSE103442 [51].

## Code availability

All codes are fully open source and available via GitHub at https://github.com/jcsia/AtBotML.

## Acknowledgements

The authors wish to thank Dr. Kenichi Tsuda for providing original bacteria growth phenotypic data, and Sylvain Raffaele for providing comments on *S. sclerotiorum* disease index. This work is supported by the National Science Foundation (NSF) through the Models for Uncovering Rules and Unexpected Phenomena in Biological Systems (MODULUS) (award no. MCB-1936800 to DEC and MCB-1936775 to PB), United State Department of Agriculture-National Institute of Food and Agriculture (USDA-NIFA) (award no. 2018-67013-28492) to DEC, and awards to PB through the NSF Career Award (CPS/CNS-1453860), the NSF awards (CCF-1837131, CNS-1932620, and CMMI-1936624), the DARPA Young Faculty Award and Director’s Fellowship Award (N66001-17-1-4044), and a Northrop Grumman grant. The funders had no role in study design, data collection and analysis, decision to publish, or preparation of the manuscript. The views, opinions, and/or findings contained in this article are those of the authors and should not be interpreted as representing the official views or policies, either expressed or implied by the Defense Advanced Research Projects Agency.

## Author contributions

All the authors contributed to the development of the project. JS and MC were responsible for data pre-processing and model construction. JS and WZ conducted data analysis with contributions by MC, PB, and DEC. All authors contributed to data interpretation and writing the manuscript.

## Supplemental Figures

**Supplemental Figure 1.**
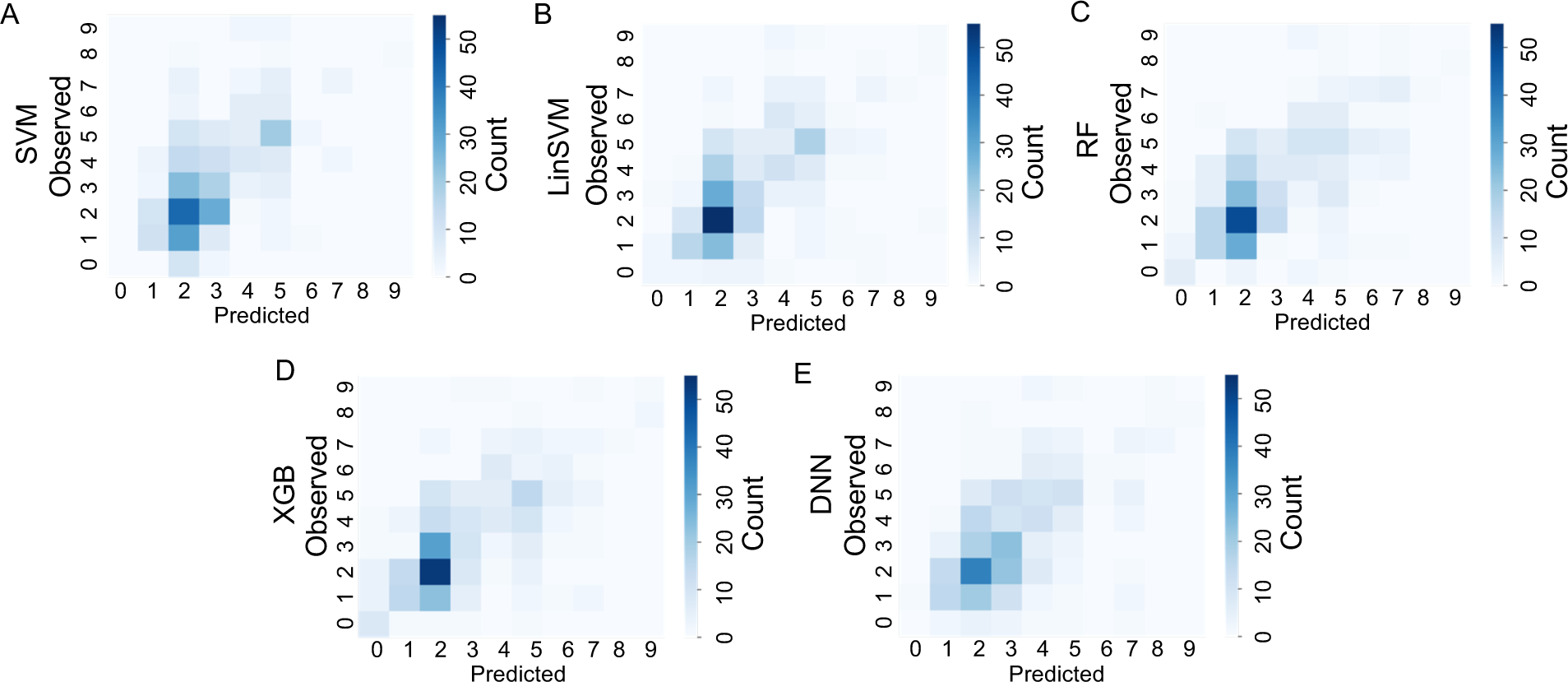
Heatmap of the observed versus predicted disease classes by machine learning model for dual-species transcriptome data. (A-E) The predicted disease score or class (x-axis), on a scale of 0 to 9, is shown versus the observed disease score (y-axis) as a heatmap. Samples where the predicted and observed values match count for a box on the diagonal axis. The total number of samples with the different predicted-observed scores are depicted in bins with the count range shown to the right of each plot. The type of ML model used is shown at the left.

**Supplemental Figure 2.**
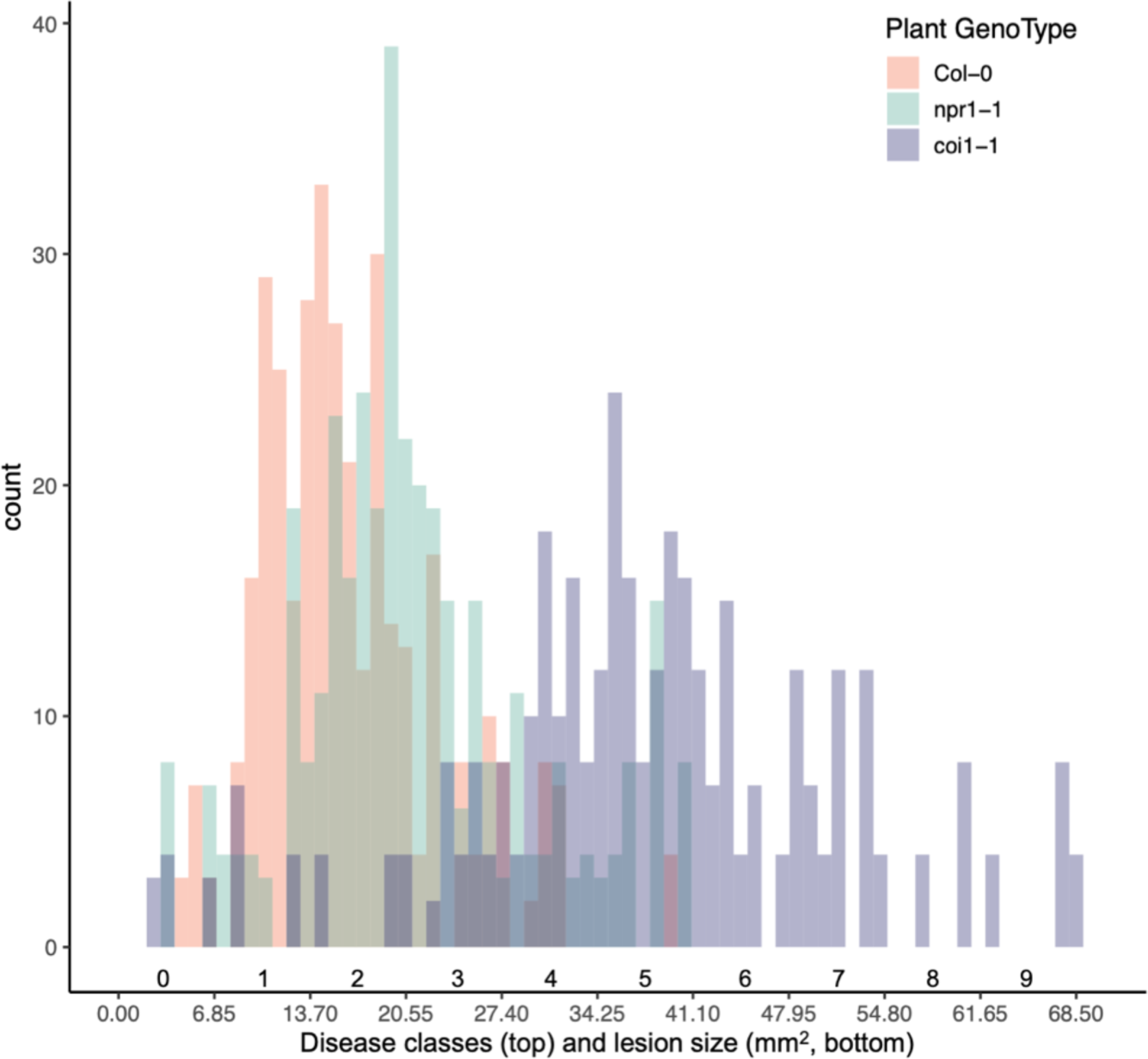
Distribution of lesion sizes during *Arabidopsis-Botrytis* interaction. The resulting lesion size (mm^2^) on *A. thaliana* leaves resulting from *B. cinerea* infection was previously collected [20]. The distribution of lesion sizes was influenced by the host genotype, color coded as shown, where *coi1* plants had larger lesion sizes. To standardize the physical lesion size into a class score that could be used across other systems, the distribution of data were fitted to a 0 to 9 scale as shown on the x-axis above the lesion size scale.

**Supplemental Figure 3.**
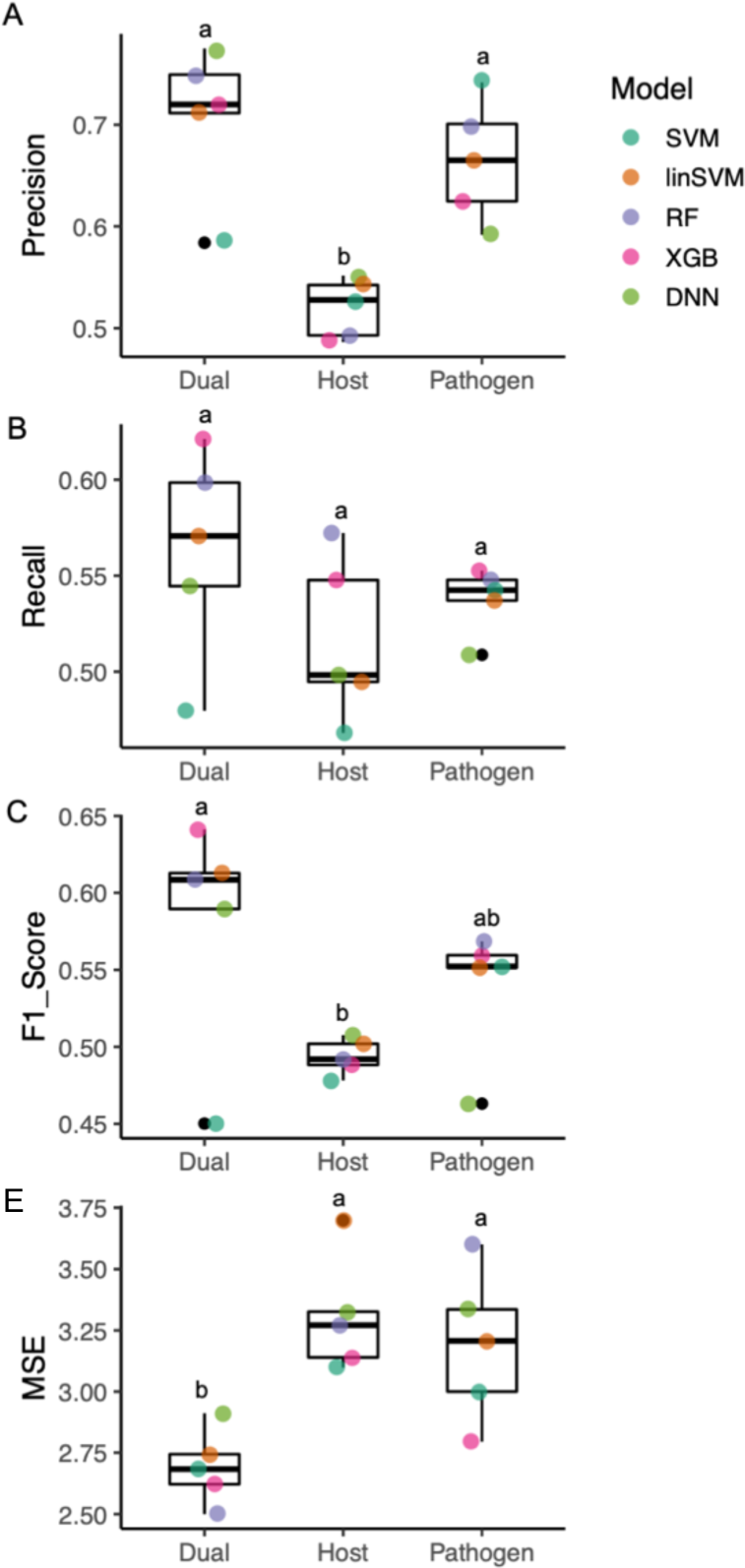
Evaluation of machine learning models based on host, fungal or dual-species transcriptomes derived from *Arabidopsis-Botrytis* interaction. Evaluation of machine learning models based on predictions of disease class ratings from *Arabidopsis-Botrytis* interaction. (A) precision (B) recall (C) F1 score (E) mean square error (MSE). For each model (A-E) the prediction was based on either both host and pathogen transcriptome data (Dual), or only *Arabidopsis* (Host) or only *Botrytis* (Pathogen). Each of the five ML models were used to test each combination of data shown as different colored points on the graphs. The letters above box blots indicate the significant levels (One-way ANOVA, p<0.05).

**Supplemental Figure 4.**
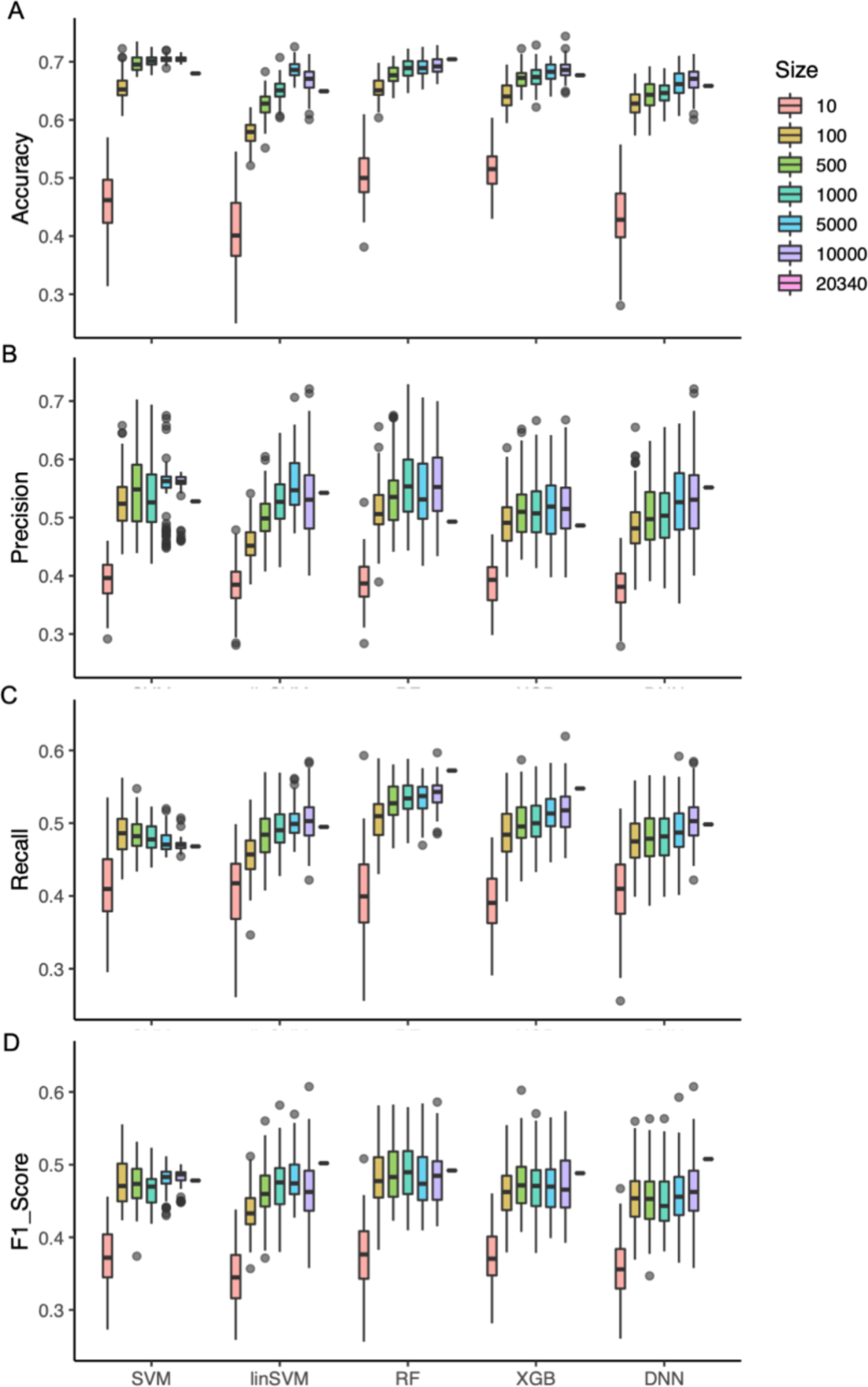
Changing the size of the feature set impacts ML performance. (A) accuracy, (B) precision, (C) recall, and (D) F1 score of predicted plant disease class by five ML models. For each model, the plant transcriptome feature size (i.e. number of genes) ranging from 10 to 20340 (the whole-genome transcriptome) were tested. The five ML models are support vector machine (SVM), linear SVM (linSVM), Random Forest (RF), XG Boost (XGB), and deep neural network (DNN).

**Supplemental Figure 5.**
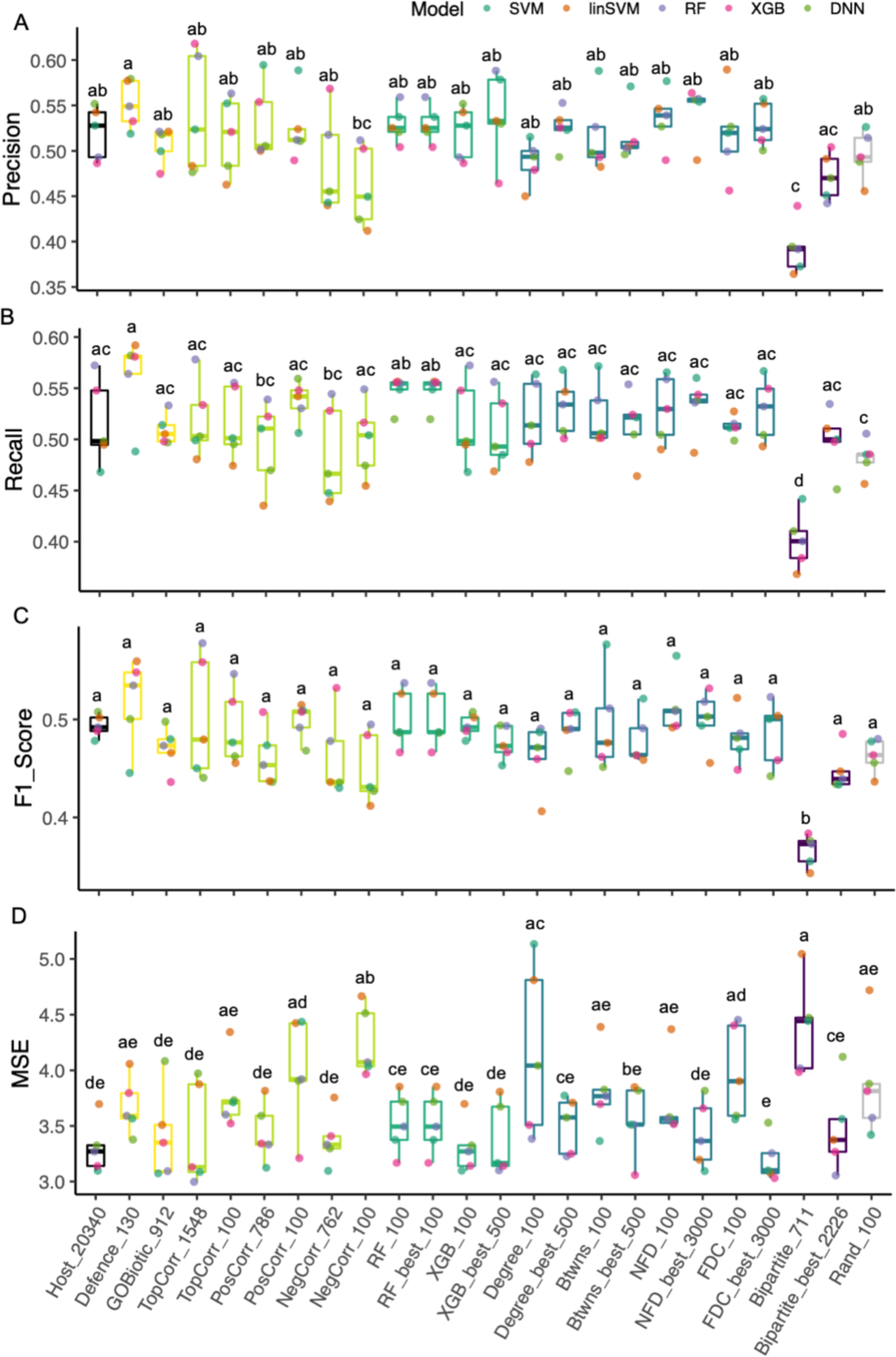
Evaluation of set size and feature selection methods impacting prediction performance during *Arabidopsis-Botrytis* interaction. (A) precision, (B) recall, (C), and (D) MSE evaluation of predicted plant disease class by each of five ML models. The five ML models were previously indicated and shown as colored points as indicated by the key at the top of the figure. For each of the feature selection methods employed here, the results for two set sizes is shown along the x-axis. The number at the end of each label is the corresponding set size. The letters above box blots indicate the significant levels (One-way ANOVA, p<0.05).

**Supplemental Figure 6.**
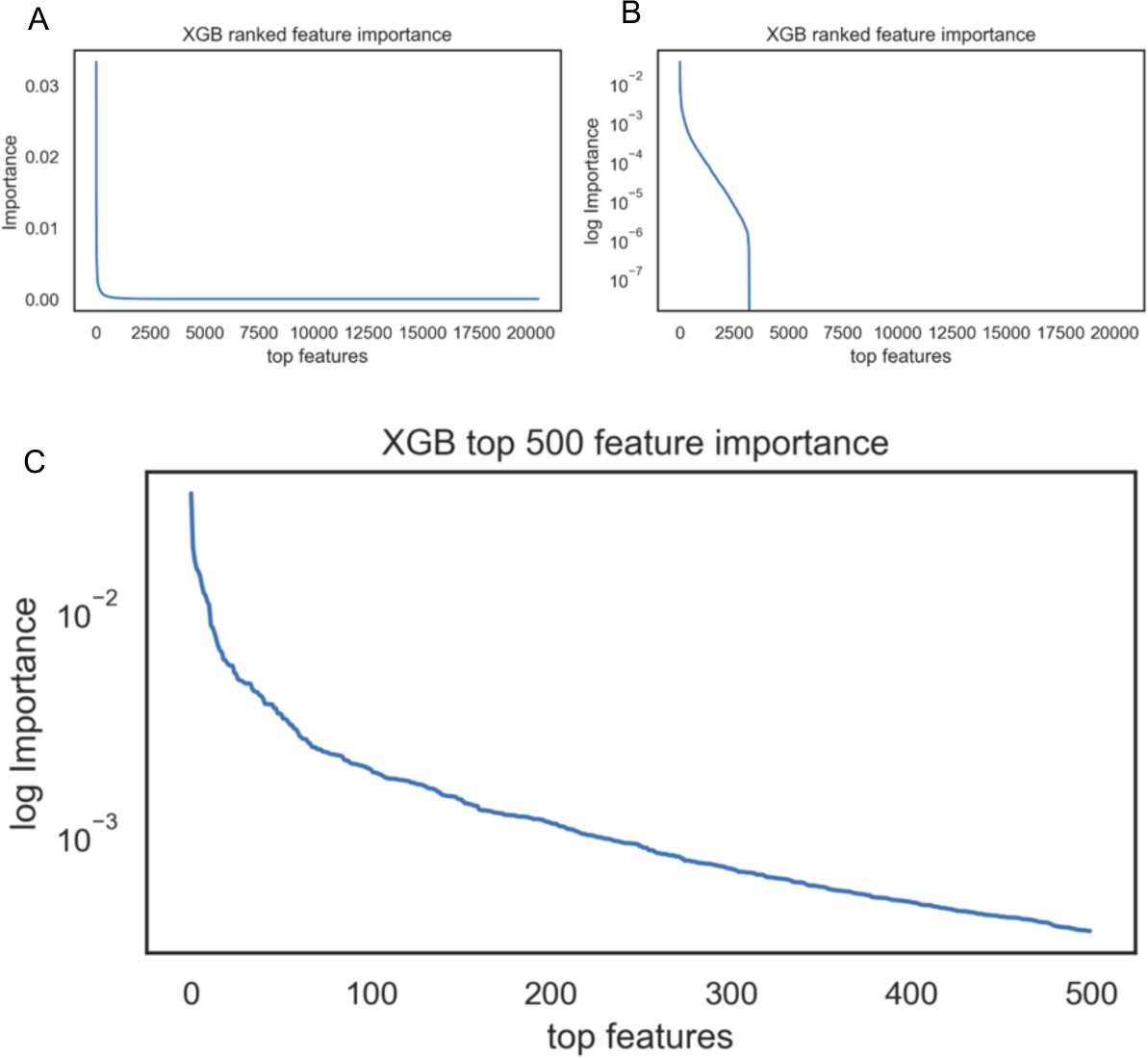
Set size and feature selection threshold selection. To determine the set size for each feature selection technique, set size was plotted by importance or performance. For example, XGBoost feature importance is illustrated for example. The feature importance is plotted on a (A) linear and (B) log scale with the set size on the x-axis. To determine which set size to use, the set size cutoff is selected at the first approximate inflection point of the log-importance curve. (C) For illustrative purposes, the log-importance for the top 500 (set size) are shown.

**Supplemental Figure 7.**
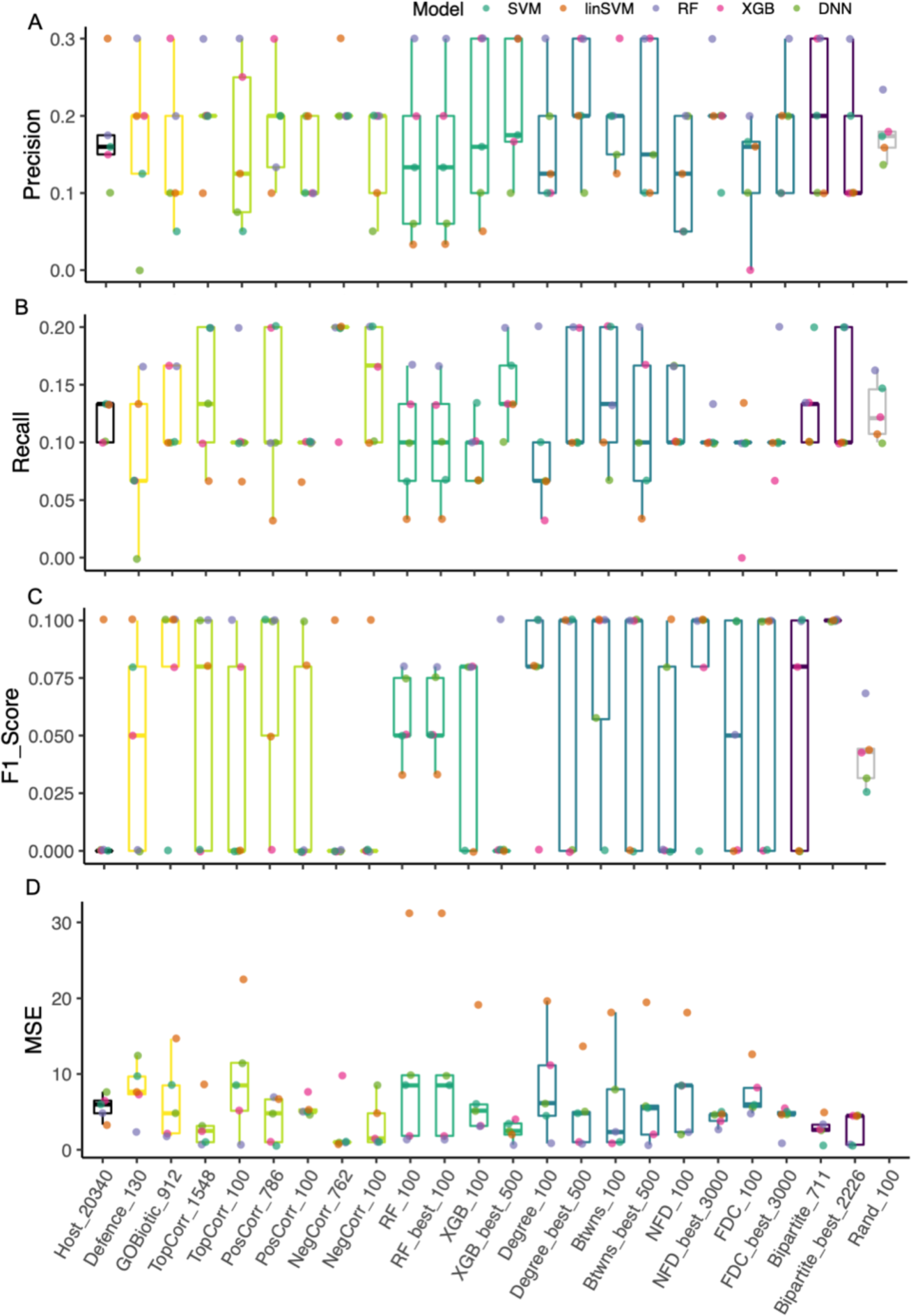
Assessment of ML models and feature selection methods on a new data set collected from *Arabidopsis*-*S. sclerotiorum* interaction. (A) precision, (B) recall, (C) F1 score, and (D) MSE evaluation of predicted plant disease class by each of five ML models. The five ML models were previously indicated and shown as colored points as indicated by the key at the top of the figure. For each of the feature selection methods employed here, the results for two set sizes is shown along the x-axis. The number at the end of each label is the corresponding set size. The letters above box blots indicate the significant levels (One-way ANOVA, p<0.05).

**Supplemental Figure 8.**
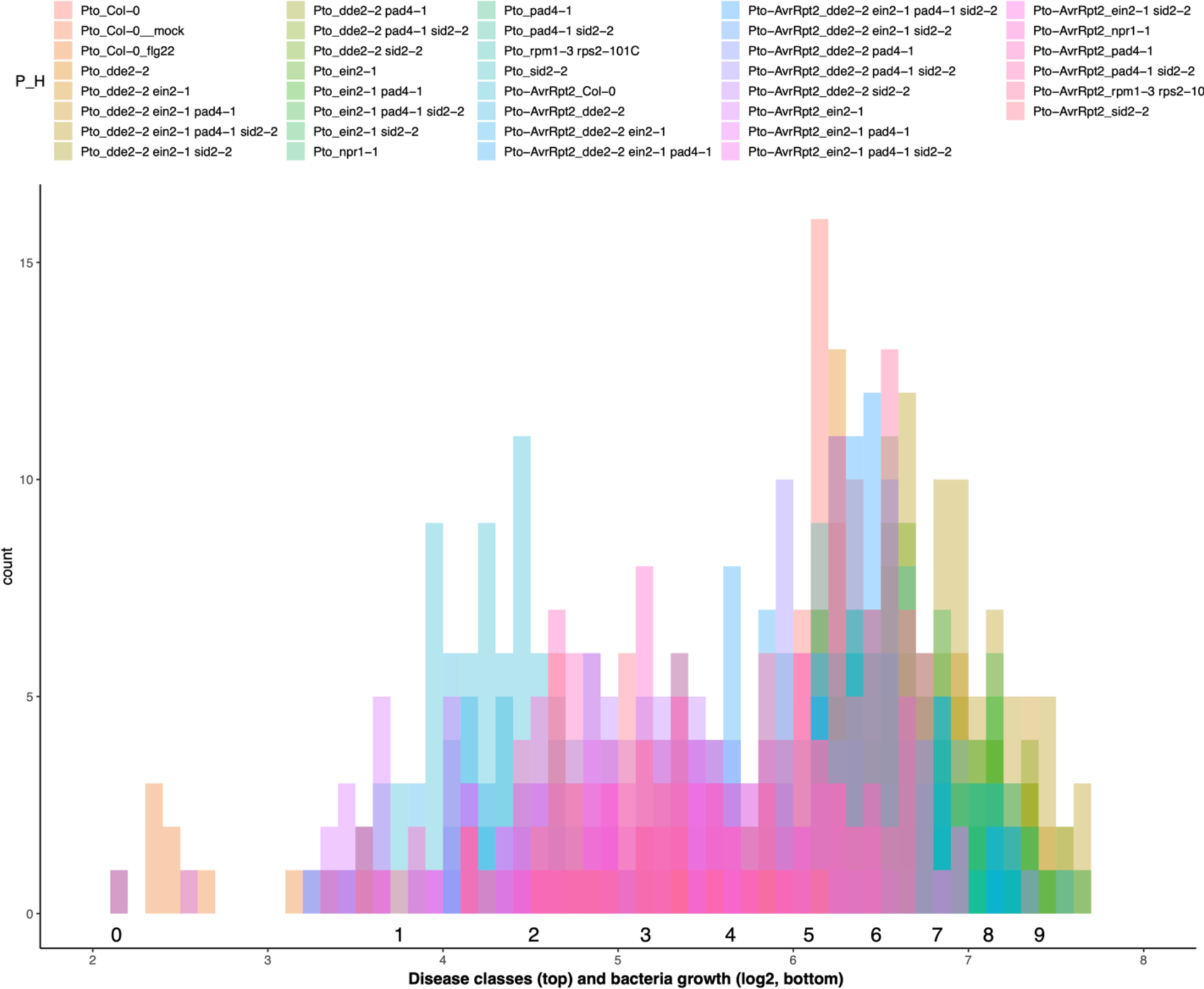
Distribution of bacteria growth derived from *Arabidopsis*-*P. syringae* interaction. The resulting bacterial growth levels (log2) from *A. thaliana* leaves infected with *P. syringae* was previously collected [51]. The distribution of bacterial growth *in planta* was influenced by the host and pathogen genotype, which is color coded for each treatment above the plot. The bacterial growth was converted to our 0 to 9 scale in order to interpret results across experiments as shown above the bacterial growth scale.

**Supplemental Figure 9.**
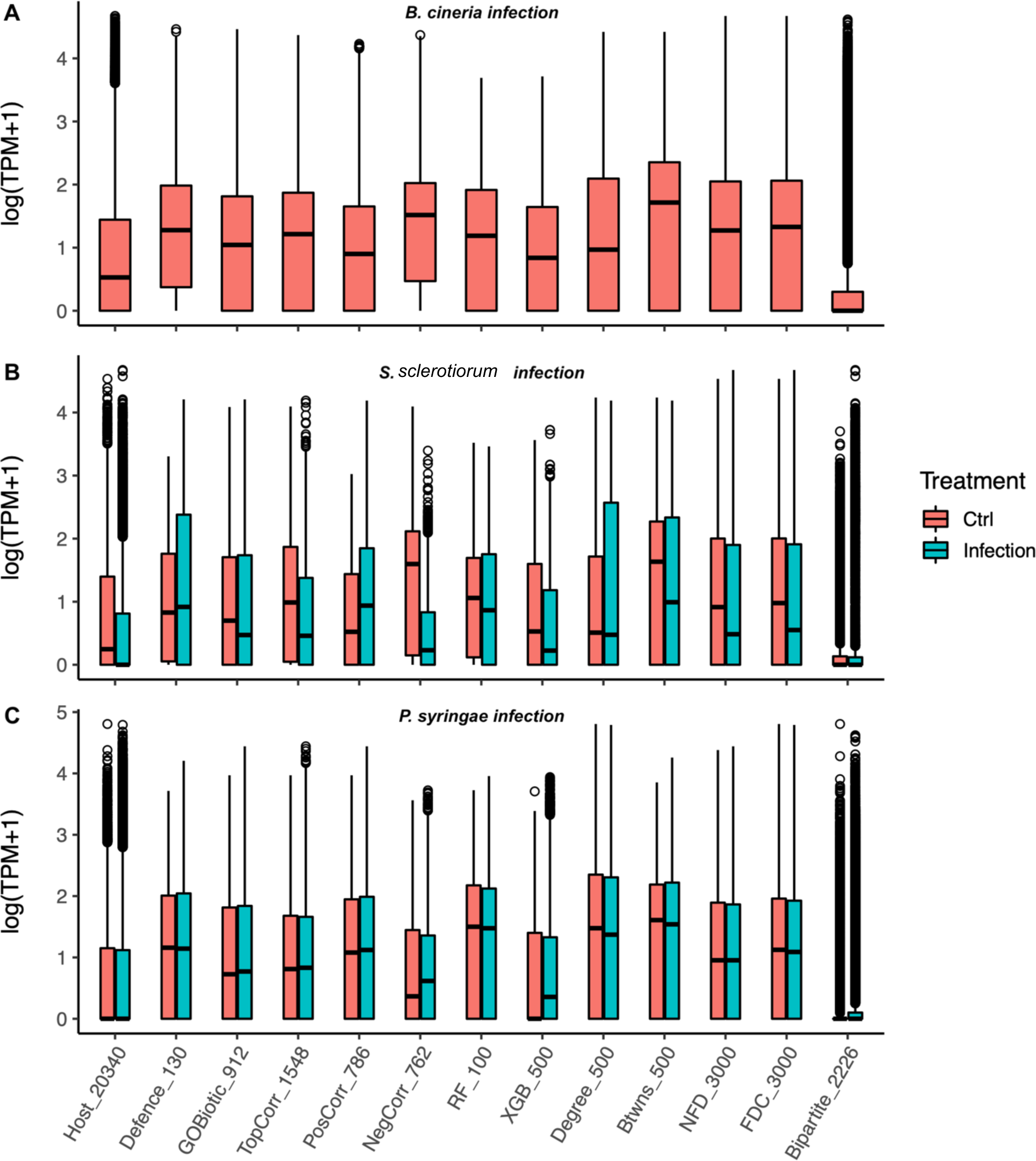
Transcript quantification for feature selection sets. plotted at the log2 value of transcripts per million plus one for (A) *B. cinerea* infection, (B) *S. sclerotiorum* infection, and (C) *P. syringae* infection. For the *S. sclerotiorum* and *P. syringae* datasets, transcript values for the feature gene sets are split between mock-inoculated and infected samples.

**Table S1.**
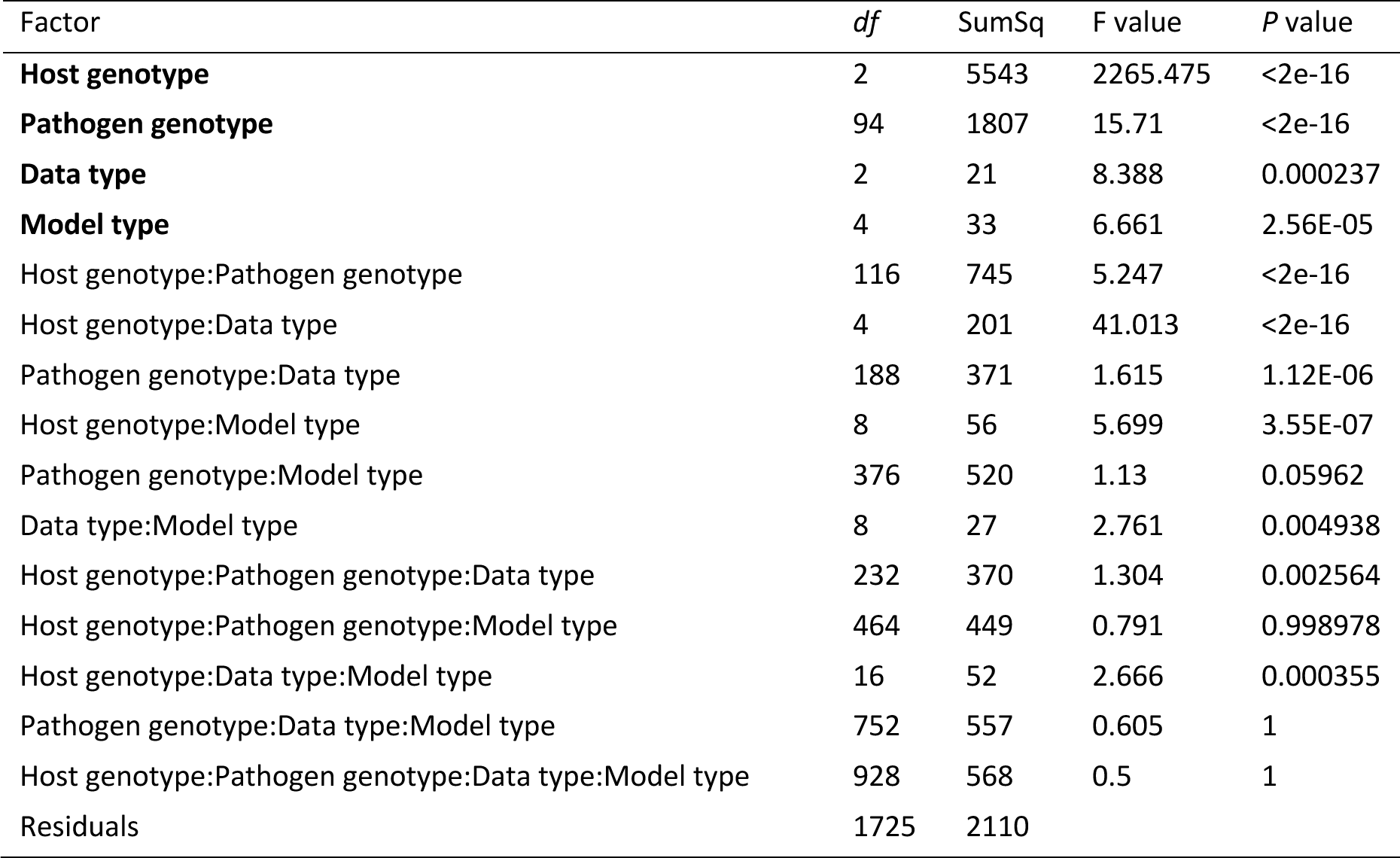
Results from ANOVA for predicted disease class errors of *Arabidopsis-Botrytis* infection.

**Table S2.** One-way ANOVA for evaluation parameters by transcriptome data types of *Arabidopsis-Botrytis* infection.

| CM parameter | Factor | *df* | SumSq | F value | *P* value |
| --- | --- | --- | --- | --- | --- |
| Accuracy | Data type | 2 | 0.004842 | 6.129 | 0.0147 |
|  | Residuals | 12 | 0.00474 |  |  |
| Precision | Data type | 2 | 0.09653 | 14.72 | 0.000589 |
|  | Residuals | 12 | 0.03934 |  |  |
| Recall | Data type | 2 | 0.005469 | 1.607 | 0.241 |
|  | Residuals | 12 | 0.020418 |  |  |
| F1 Score | Data type | 2 | 0.01886 | 3.712 | 0.0556 |
|  | Residuals | 12 | 0.03049 |  |  |
| Mean Square Error | Data type | 2 | 1.0628 | 9.101 | 0.00393 |
|  | Residuals | 12 | 0.7006 |  |  |

**Table S3.**
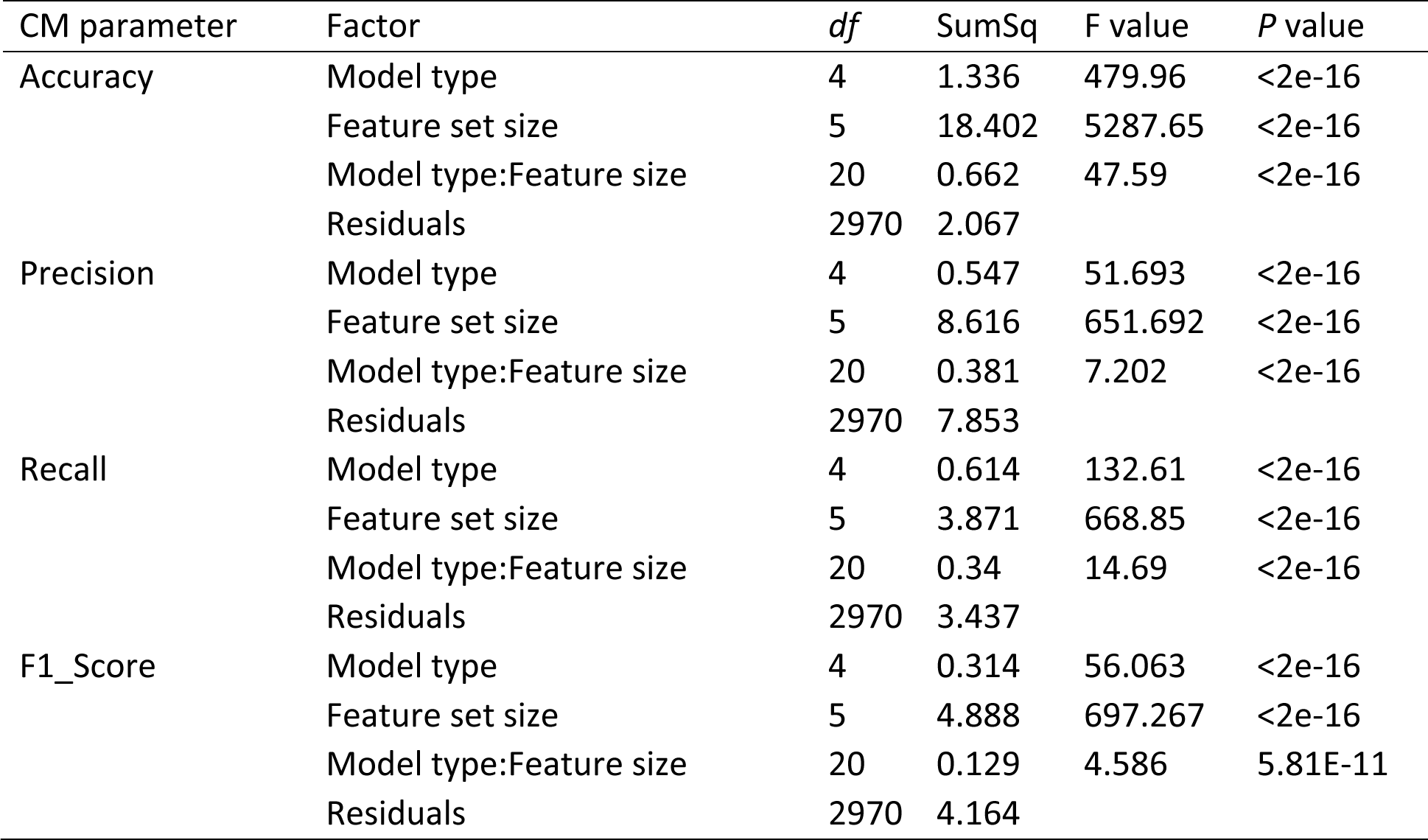
Two-way ANOVA on evaluation parameters by effects of machine learning models and feature size of *Arabidopsis-Botrytis* infection.

**Table S4. Gene lists derived from diverse feature selection methods.** Available as an excel file table due to size

**Table S5.** Two-way ANOVA for performance parameters by effects of feature selection methods and ML models on *Arabidopsis-Botrytis* pathosystem.

| CM parameter | Factor | *df* | SumSq | F value | *P* value |
| --- | --- | --- | --- | --- | --- |
| Accuracy | Feature selection method | 27 | 0.4762 | 25.98 | < 2.00e-16 |
|  | ML model type | 4 | 0.0643 | 23.68 | 4.46E-14 |
|  | Residuals | 108 | 0.0733 |  |  |
| Precision | Feature selection method | 27 | 0.25048 | 7.926 | 1.84E-15 |
|  | ML model type | 4 | 0.01202 | 2.568 | 4.21E-02 |
|  | Residuals | 108 | 0.1264 |  |  |
| Recall | Feature selection method | 27 | 0.1446 | 6.249 | 2.33E-12 |
|  | ML model type | 4 | 0.03078 | 8.978 | 2.68E-06 |
|  | Residuals | 108 | 0.09256 |  |  |
| F1 Score | Feature selection method | 27 | 0.20963 | 8.37 | 3.20E-16 |
|  | ML model type | 4 | 0.01666 | 4.49 | 2.15E-03 |
|  | Residuals | 108 | 0.10019 |  |  |

**Table S6.**
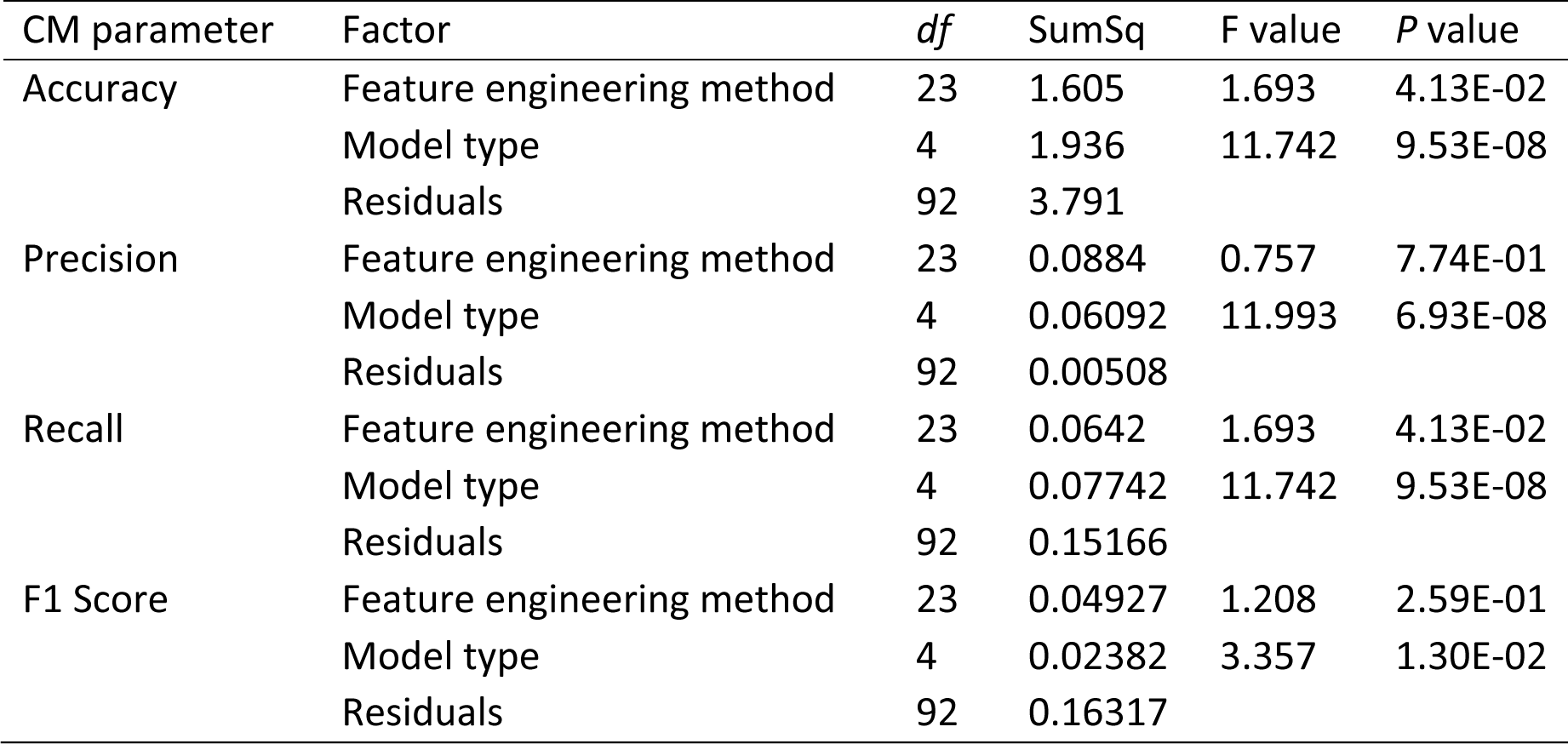
Two-way ANOVA on five evaluation parameters by effects of feature selection methods and machine learning models on *Arabidopsis-S.sclerotiorum* pathosystem.

| CM parameter | Factor | *df* | SumSq | F value | *P* value |
| --- | --- | --- | --- | --- | --- |
| Accuracy | Feature engineering method | 23 | 1.605 | 1.693 | 4.13E-02 |
|  | Model type | 4 | 1.936 | 11.742 | 9.53E-08 |
|  | Residuals | 92 | 3.791 |  |  |
| Precision | Feature engineering method | 23 | 0.0884 | 0.757 | 7.74E-01 |
|  | Model type | 4 | 0.06092 | 11.993 | 6.93E-08 |
|  | Residuals | 92 | 0.00508 |  |  |
| Recall | Feature engineering method | 23 | 0.0642 | 1.693 | 4.13E-02 |
|  | Model type | 4 | 0.07742 | 11.742 | 9.53E-08 |
|  | Residuals | 92 | 0.15166 |  |  |
| F1 Score | Feature engineering method | 23 | 0.04927 | 1.208 | 2.59E-01 |
|  | Model type | 4 | 0.02382 | 3.357 | 1.30E-02 |
|  | Residuals | 92 | 0.16317 |  |  |

**Table S7.** Results ANOVA for predicted disease class error of *A.thaliana-P.syringae* infection.

| Factor | *df* | SumSq | F value | *P* value |
| --- | --- | --- | --- | --- |
| Host Immunity | 4 | 9156 | 1023.63 | <2e-16 |
| Pathogen genotype | 1 | 28529 | 12758.45 | <2e-16 |
| Model type | 4 | 299 | 33.42 | <2e-16 |
| Feature engineering method | 22 | 7032 | 142.94 | <2e-16 |
| Host Immunity:Pathogen genotype | 3 | 1054 | 157.13 | <2e-16 |
| Host Immunity:Model type | 16 | 228 | 6.36 | 2.11E-14 |
| Pathogen genotype:Model type | 4 | 271 | 30.33 | <2e-16 |
| Host Immunity:Feature engineering method | 88 | 476 | 2.42 | 3.99E-12 |
| Pathogen genotype:Feature engineering method | 22 | 419 | 8.52 | <2e-16 |
| Model type:Feature engineering method | 69 | 11326 | 73.41 | <2e-16 |
| Host Immunity:Pathogen genotype:Model type | 12 | 40 | 1.50 | 0.1164 |
| Host Immunity:Pathogen genotype:Feature engineering method | 66 | 208 | 1.41 | 0.0165 |
| Host Immunity:Model type:Feature engineering method | 276 | 1084 | 1.76 | 5.29E-13 |
| Pathogen genotype:Model type:Feature engineering method | 69 | 1132 | 7.33 | <2e-16 |
| Host Immunity:Pathogen genotype:Model type:Feature engineering method | 207 | 546 | 1.18 | 0.0416 |
| Residuals | 7104 | 15885 |  |  |

**Table S8.**
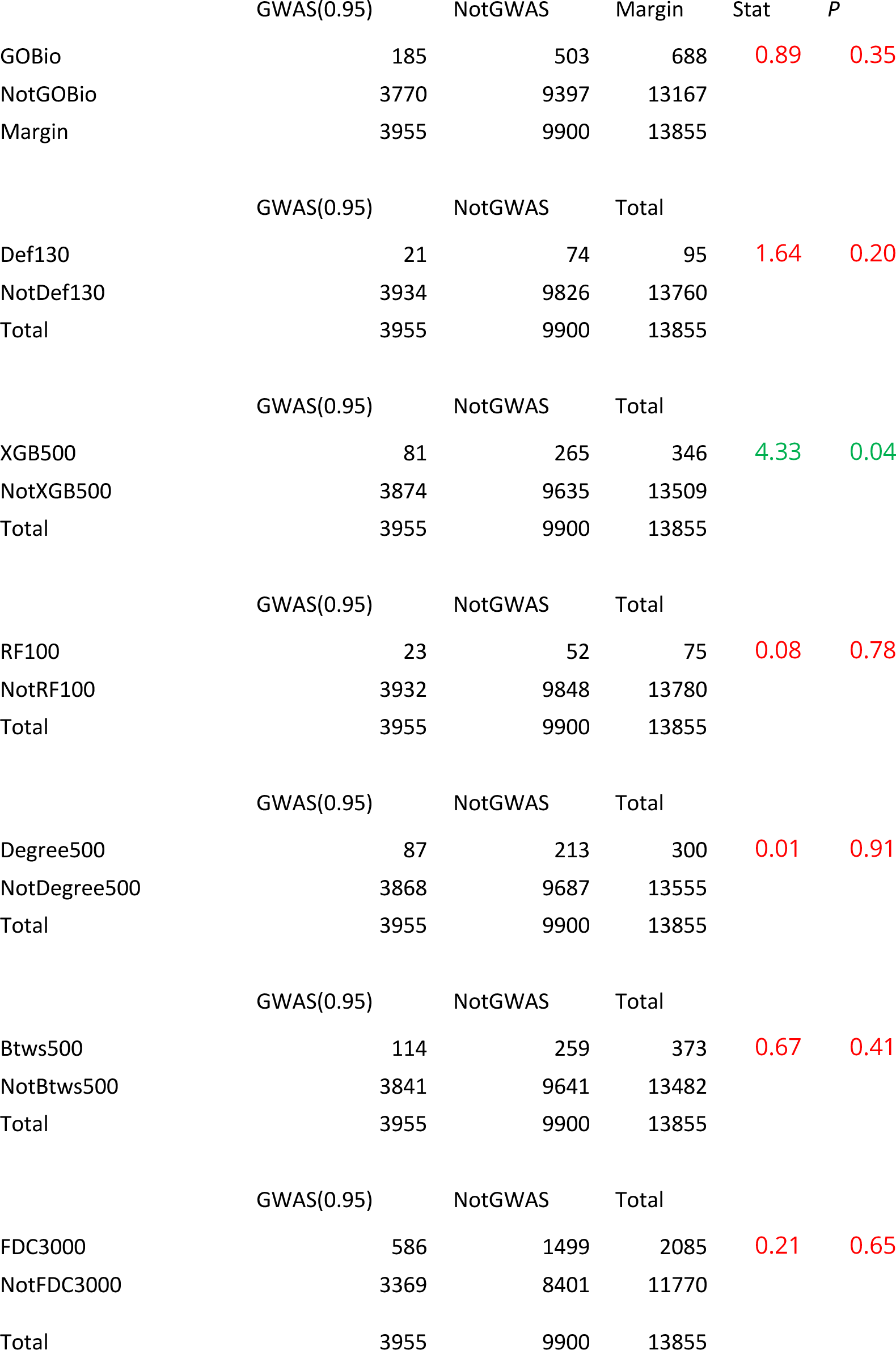
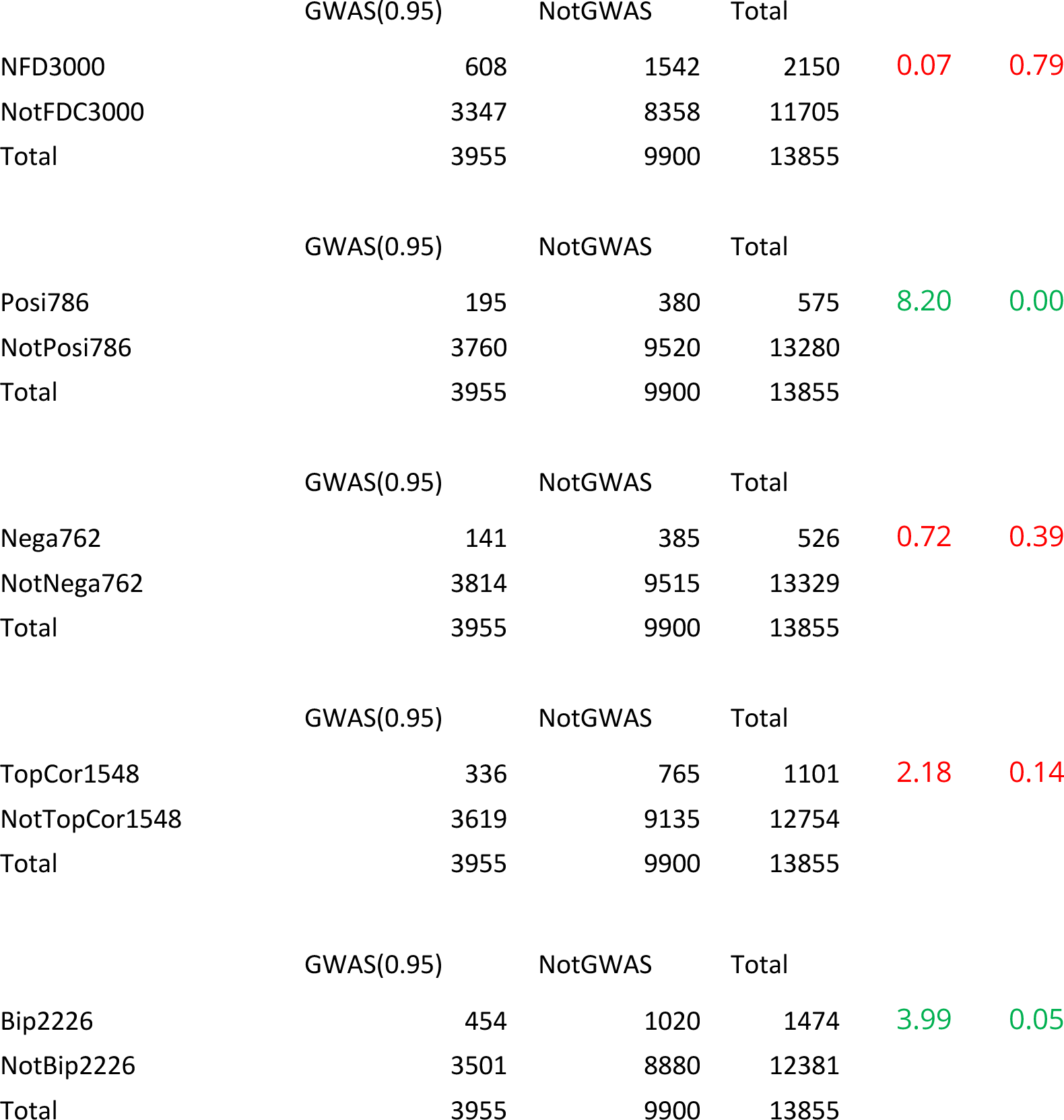
Chi-square results for GWAS gene set with each of the feature selection sets.

|  | GWAS(0.95) | NotGWAS | Margin | Stat | P |
| --- | --- | --- | --- | --- | --- |
| GOBio | 185 | 503 | 688 | 0.89 | 0.35 |
| NotGOBio | 3770 | 9397 | 13167 |  |  |
| Margin | 3955 | 9900 | 13855 |  |  |
|  | GWAS(0.95) | NotGWAS | Total |  |  |
| Def130 | 21 | 74 | 95 | 1.64 | 0.20 |
| NotDef130 | 3934 | 9826 | 13760 |  |  |
| Total | 3955 | 9900 | 13855 |  |  |
|  | GWAS(0.95) | NotGWAS | Total |  |  |
| XGB500 | 81 | 265 | 346 | 4.33 | 0.04 |
| NotXGB500 | 3874 | 9635 | 13509 |  |  |
| Total | 3955 | 9900 | 13855 |  |  |
|  | GWAS(0.95) | NotGWAS | Total |  |  |
| RF100 | 23 | 52 | 75 | 0.08 | 0.78 |
| NotRF100 | 3932 | 9848 | 13780 |  |  |
| Total | 3955 | 9900 | 13855 |  |  |
|  | GWAS(0.95) | NotGWAS | Total |  |  |
| Degree500 | 87 | 213 | 300 | 0.01 | 0.91 |
| NotDegree500 | 3868 | 9687 | 13555 |  |  |
| Total | 3955 | 9900 | 13855 |  |  |
|  | GWAS(0.95) | NotGWAS | Total |  |  |
| Btws500 | 114 | 259 | 373 | 0.67 | 0.41 |
| NotBtws500 | 3841 | 9641 | 13482 |  |  |
| Total | 3955 | 9900 | 13855 |  |  |
|  | GWAS(0.95) | NotGWAS | Total |  |  |
| FDC3000 | 586 | 1499 | 2085 | 0.21 | 0.65 |
| NotFDC3000 | 3369 | 8401 | 11770 |  |  |
| Total | 3955 | 9900 | 13855 |  |  |

|  | GWAS(0.95) | NotGWAS | Total |  |  |
| --- | --- | --- | --- | --- | --- |
| NFD3000 | 608 | 1542 | 2150 | 0.07 | 0.79 |
| NotFDC3000 | 3347 | 8358 | 11705 |  |  |
| Total | 3955 | 9900 | 13855 |  |  |
| Posi786 | 195 | 380 | 575 | 8.20 | 0.00 |
| NotPosi786 | 3760 | 9520 | 13280 |  |  |
| Total | 3955 | 9900 | 13855 |  |  |
| Nega762 | 141 | 385 | 526 | 0.72 | 0.39 |
| NotNega762 | 3814 | 9515 | 13329 |  |  |
| Total | 3955 | 9900 | 13855 |  |  |
| TopCor1548 | 336 | 765 | 1101 | 2.18 | 0.14 |
| NotTopCor1548 | 3619 | 9135 | 12754 |  |  |
| Total | 3955 | 9900 | 13855 |  |  |
| Bip2226 | 454 | 1020 | 1474 | 3.99 | 0.05 |
| NotBip2226 | 3501 | 8880 | 12381 |  |  |
| Total | 3955 | 9900 | 13855 |  |  |

**Table S9.** Chi-square results for Bjornson identified gene set with each of the feature selection sets.

|  | Bjornson | NotBjornson | Margin | Stat | P |
| --- | --- | --- | --- | --- | --- |
| GOBio | 186 | 726 | 912 | 509.74 | <0.0001 |
| NotGOBio | 784 | 18644 | 19428 |  |  |
| Total | 970 | 19370 | 20340 |  |  |
|  | Bjornson | NotBjornson | Margin |  |  |
| Def130 | 36 | 94 | 130 | 146.35 | <0.0001 |
| NotDef130 | 934 | 19276 | 20210 |  |  |
| Total | 970 | 19370 | 20340 |  |  |
|  | Bjornson | NotBjornson | Margin |  |  |
| XGB500 | 20 | 480 | 500 | 0.51 | 0.48 |
| NotXGB500 | 950 | 18890 | 19840 |  |  |
| Total | 970 | 19370 | 20340 |  |  |
|  | Bjornson | NotBjornson | Margin |  |  |
| RF100 | 24 | 76 | 100 | 77.64 | <0.0001 |
| NotRF100 | 946 | 19294 | 20240 |  |  |
| Total | 970 | 19370 | 20340 |  |  |
|  | Bjornson | NotBjornson | Margin |  |  |
| Degree500 | 75 | 425 | 500 | 115.85 | <0.0001 |
| NotDegree500 | 895 | 18945 | 19840 |  |  |
| Total | 970 | 19370 | 20340 |  |  |
|  | Bjornson | NotBjornson | Margin |  |  |
| Btws500 | 82 | 418 | 500 | 150.08 | <0.0001 |
| NotBtws500 | 888 | 18952 | 19840 |  |  |
| Total | 970 | 19370 | 20340 |  |  |
|  | Bjornson | NotBjornson | Margin |  |  |
| FDC3000 | 340 | 2660 | 3000 | 332.21 | <0.0001 |
| NotFDC3000 | 630 | 16710 | 17340 |  |  |
| Total | 970 | 19370 | 20340 |  |  |
|  | Bjornson | NotBjornson | Margin |  |  |
| NFD3000 | 335 | 2665 | 3000 | 315.51 | <0.0001 |
| NotNFD3000 | 635 | 16705 | 17340 |  |  |
| Total | 970 | 19370 | 20340 |  |  |
|  | Bjornson | NotBjornson | Margin |  |  |
| Posi786 | 257 | 529 | 786 | 1397.80 | <0.0001 |
| NotPosi786 | 713 | 18841 | 19554 |  |  |
| Total | 970 | 19370 | 20340 |  |  |
|  | Bjornson | NotBjornson | Margin |  |  |
| Nega762 | 0 | 762 | 762 | 38.56 | <0.0001 |
| NotNega762 | 970 | 18608 | 19578 |  |  |
| Total | 970 | 19370 | 20340 |  |  |
|  | Bjornson | NotBjornson | Margin |  |  |
| TopCor1548 | 257 | 1291 | 1548 | 513.78 | <0.0001 |
| NotTopCor1548 | 713 | 18079 | 18792 |  |  |
| Total | 970 | 19370 | 20340 |  |  |
|  | Bjornson | NotBjornson | Margin |  |  |
| Bip2226 | 98 | 2128 | 2226 | 0.65 | 0.42 |
| NotBip2226 | 872 | 17242 | 18114 |  |  |
| Total | 970 | 19370 | 20340 |  |  |

